# Essential sterols from engineered yeast prevent honeybee colony population decline

**DOI:** 10.1101/2024.12.09.627483

**Authors:** Elynor Moore, Raquel T de Sousa, Stella Felsinger, Jonathan A Arnesen, Jane D Dyekjær, Dudley I Farman, Rui F S Gonçalves, Philip C Stevenson, Irina Borodina, Geraldine A Wright

**Affiliations:** Department of Biology, University of Oxford, Oxford, United Kingdom; The Novo Nordisk Foundation Centre for Biosustainability, Technical University of Denmark, Kongens Lyngby, Denmark; Natural Resources Institute, University of Greenwich, Chatham, United Kingdom; Kew Science Trait Diversity and Function, Royal Botanic Gardens Kew, Richmond, United Kingdom

## Abstract

Honeybees, the world’s most important crop pollinators, are increasingly facing pollen starvation arising from agricultural intensification and climate change. Frequent flowering dearth periods and high-density rearing conditions weaken colonies, often leading to their demise. Beekeepers provide colonies with pollen substitutes, but these feeds cannot sustain brood production because they lack essential sterols found in pollen. Here, we describe a technological breakthrough in honeybee nutrition with wide-reaching impacts on global food security. We first measured the quantity and proportion of sterols found in honeybee tissues. Using this information, we genetically engineered a strain of the oleaginous yeast, *Yarrowia lipolytica*, to produce a mixture of essential sterols for bees and incorporated it into an otherwise nutritionally complete diet. Colonies fed exclusively with this diet reared brood for significantly longer than those fed diets without suitable sterols. Incorporating sterol supplements into pollen substitutes using this method will enable beekeepers to rear healthier, longer-lived colonies to meet the growing demands for global crop pollination. It could also reduce competition between bee species for access to natural floral resources, stemming the decline of wild bee populations.

## Main

Managed honeybees (*Apis mellifera*) are fundamental to modern agricultural systems and global food security due to the pollination services they provide to crops. Honeybees consume floral pollen, which provides protein, fats, carbohydrate, and micronutrients^1^. Increasingly, a combination of changes in climate, land use, and agricultural practises leaves honeybees with limited access to sufficient and diverse floral resources^2–4^. Nutritional deficiencies increase susceptibility to disease and colony collapse, thereby contributing to the growing rate of honeybee colony losses^5,6^. Loss of pollinators, in turn, reduces crop yields and increases the costs of food production^7,8^. In the past 40 years, beekeepers have adopted the practice of feeding artificial pollen substitutes where natural forage is insufficient, or bees are kept at high densities. However, commercially available pollen substitutes composed of protein flour, sugars, and oils are not nutritionally complete feeds for honeybees. They are missing essential sterols found in floral pollen which are necessary for colony health and growth^9,10^.

Sterols are a structurally diverse class of tetracyclic triterpenoids that are ubiquitously important for eukaryotic cell function, notably for membrane architecture, hormone biosynthesis and signalling^11–13^. Unlike other animals, insects and marine invertebrates have lost the ability to synthesise endogenous sterols, acquiring them instead from diet^14,15^. Most insects produce cholesterol (CHOL) by dealkylating dietary sterols^15,16^. In contrast, honeybees have lost the ability to dealkylate sterols and instead use dietary sterols directly as membrane inserts and as precursors for ecdysteroid hormones^17,18^. Most of the pollen sterols used by bees are not available in quantities that could be fed to bee colonies on a commercial scale, making it impossible to create a nutritionally complete feed that is a substitute for pollen.

### Honeybees acquire six key sterols from floral pollen

Sterols are consumed by nurse-aged worker honeybees from stored pollen (bee bread); these accumulate in their mandibular glands, and are transferred to worker, queen, and drone larvae via glandular secretions (Fig. 1a)^19^. Sterol diversity is high in pollen^20^, but only a few key sterols appear to be taken up by workers in the gut and transferred to the brood (Fig. 1a, b)^21^. To quantify the proportions of sterols found in honeybees, we analysed the sterol composition in pupal tissue from each caste; the delta-5 sterol, 24-methylenecholesterol (24-MC, Extended Data Fig. 1), constituted 60-70% of the corporeal sterols (Fig. 1b). Five other minor sterols were consistently present: beta-sitosterol (SITO), desmosterol (DESMO), isofucosterol (ISOFUC), campesterol (CAMP) and cholesterol (CHOL, Fig. 1b, Extended Data Fig. 1). The relative abundance of each sterol found in pupae varied among castes (two-way quasi-binomial GLM, sterol*caste, χ^2^(10) = 47.0, *P* < 0.0001, Fig. 1b). Drone pupae had 11.8% more 24-MC than worker pupae and 2.24-fold more ISOFUC than queen pupae. Worker pupae had at least 1.5-fold more DESMO than queen or drone pupae. CAMP was almost six times more abundant in worker and queen pupae than drone pupae.

**Fig. 1:**
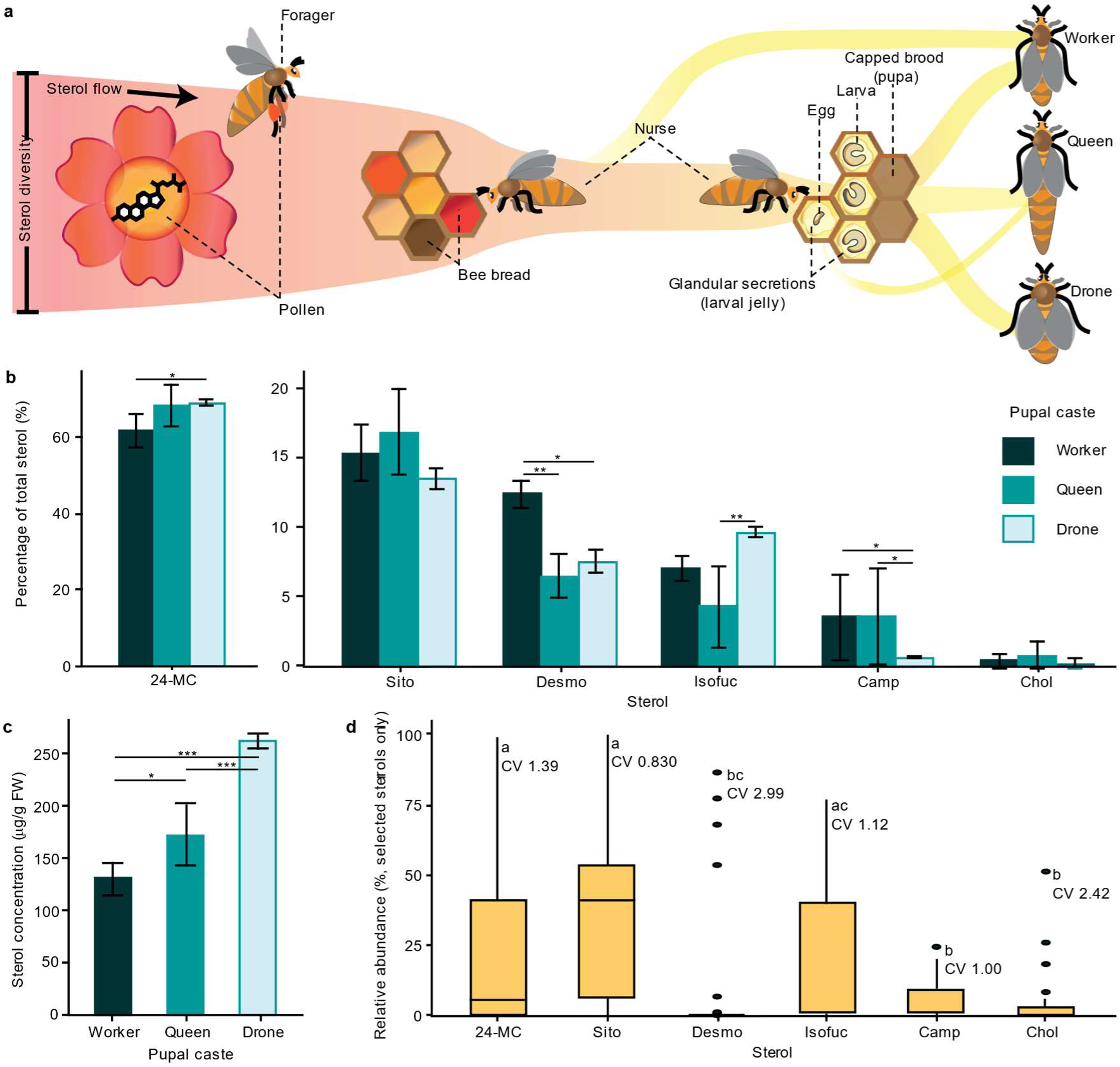
Sterol nutrition of a honeybee colony. **a**, Adult worker honeybees collect and eat floral pollen, which provides protein, lipids, and micronutrients. Pollen foraged from flowers is packed into cells of the wax comb with honey to form bee bread. Nurse bees consume bee bread and create glandular secretions (jelly) to feed larvae that include sterols required for development. Larvae develop into pupae and eventually emerge as female workers, queens, or drones (males). **b**, The relative abundance of each of the six key sterols found in pupae varies among castes (*n* = 5). **c**, Total sterol concentration varies significantly between castes (*n* = 5). **d**, Relative abundance of selected sterols in pollen from bee-pollinated flowers (% combined total, *n* = 41). Coefficients of variation (CV) of each sterol are shown. Data obtained from Zu *et al*, 2021 (Supplementary Data 1)^20^. Error bars represent ± s.d; letters and * denote statistical significance (*post hoc* comparisons of estimated marginal means with Tukey adjustment, \**P* < 0.05, \*\**P* < 0.01, \*\*\**P* < 0.001).

The total sterol concentration in pupal tissue varied significantly between castes and ranged from 130-260 μg/g fresh tissue (Gaussian GLM, χ^2^(2) = 91.9, *P* < 0.0001, Fig. 1c). Total sterol concentration in drone pupae was greater than in queen pupae and double that in worker pupae. The concentrations of individual sterols similarly varied between castes (Extended Data Fig. 2). Five of these sterols (SITO, CAMP, CHOL, ISOFUC, and 24-MC) were among the six most common sterols found in pollen (SITO, CAMP, CHOL, ISOFUC, cycloartenol and 24- MC, Supplementary Data 1)^20^. Using the data of Zu *et al,* 2021^20^, we found that floral pollen varies significantly in the relative abundance of these sterols (quasi-binomial GLM, χ^2^(5) = 61.0, *P* < 0.001). SITO had the highest median relative abundance (40.9%), followed by ISOFUC (9.50%) and 24-MC (5.84%); CAMP, CHOL and DESMO were minor sterols (Fig. 1d). The coefficient of variation ranged from 0.830 (SITO) to 2.99 (DESMO, Fig. 1d).

### *Y. lipolytica* can be engineered to produce rare pollen sterols at high content

To acquire the pollen sterols needed by bees, we adopted the oleaginous yeast, *Yarrowia lipolytica*, as a host for sterol production, due to its high capacity for lipid accumulation and abundance of the sterol precursor acetyl-CoA^22–24^. Furthermore, we used a platform strain with improved mevalonate pathway flux towards farnesyl diphosphate (FPP) for increasing sterol precursor supply^25^. The platform strain is derived from the wild-type isolate W29, express CRISPR-associated protein (Cas9) for genome editing, and overexpress four genes encoding mevalonate pathway enzymes (*HMG1*, *ERG12*, *IDI1*, *ERG20*), alongside genes involved in acetyl-CoA formation, namely ATP-citrate lyase 1 (*ACL1*) and acetyl-CoA synthase from *Salmonella enterica* (*SeACS*). Almost three times more ergosterol (ERGO, 1.95 mg/g DCW) was produced by the platform strain compared to strain W29 (0.659 mg/g DCW, Extended Data Fig. 3a).

The endogenous ERGO pathway of yeasts can be remodelled to synthesise alternative sterols (Fig. 2), but previous attempts to produce non-native sterols in *Y. lipolytica* have struggled to achieve high specific yield or stereoselectivity (Supplementary Table 1). In *S. cerevisiae*, the latter ERGO pathway genes are nonessential and alternative sterols can be taken up from the environment when ERGO biosynthesis is compromised, such as under anaerobic conditions^26,27^. However, most non-*Saccharomyces* yeasts, such as *Y. lipolytica,* lack sterol uptake transporters and do not exhibit flexible sterol auxotrophy^28^. For this reason, endogenous ERGO synthesis is essential, making deletion of the latter ERGO pathway genes such as *ERG4* (delta-24(28) sterol reductase) challenging in *Y. lipolytica*^29^.

**Fig. 2:**
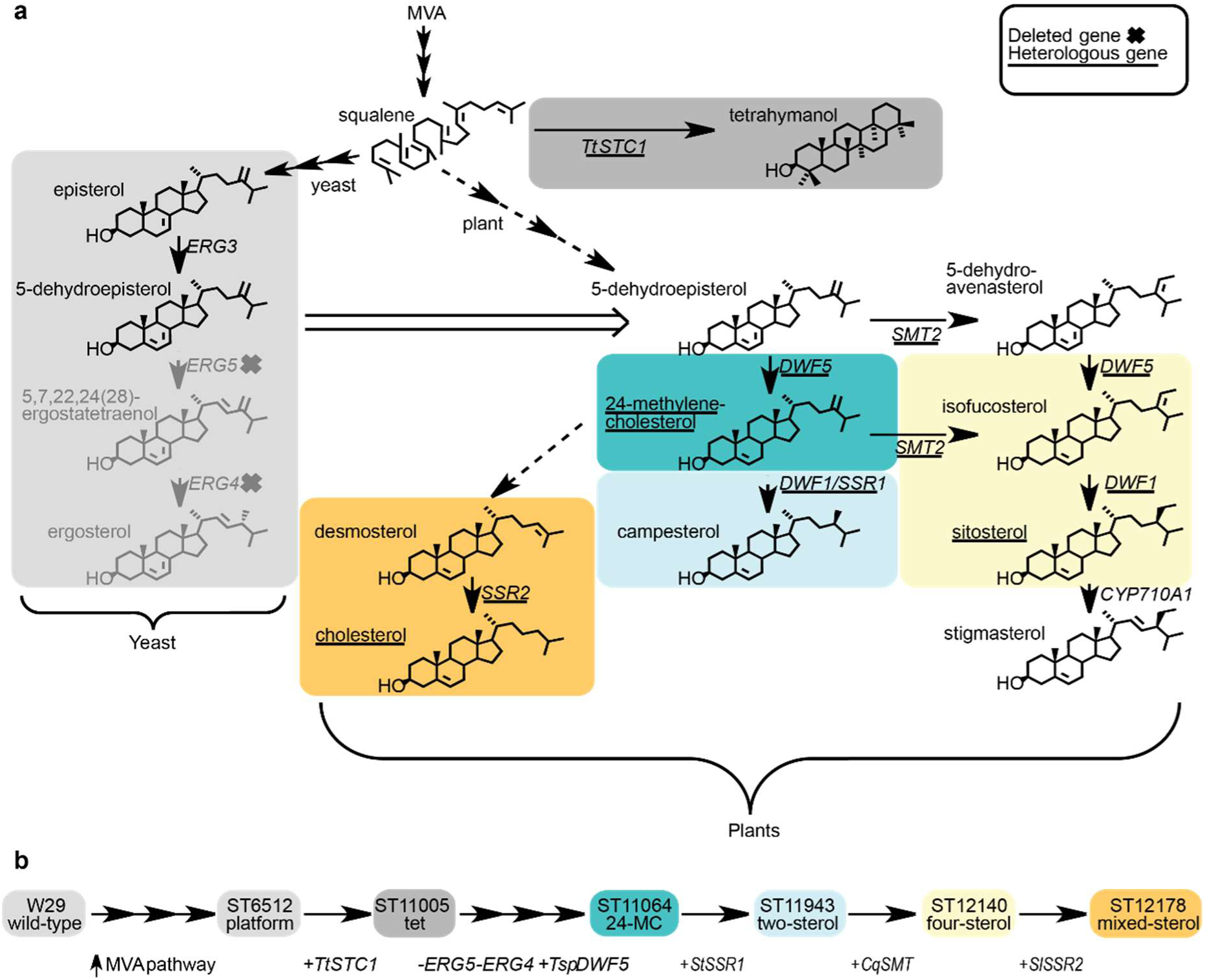
Biosynthetic pathways to produce the sterols used by bees, in plants and engineered yeast. **a**, Schematic of sterol biosynthetic pathway from mevalonate (MVA) in yeast and plants. The latter steps in the pathways and the enzymes that catalyse these reactions are indicated: *ERG3* (EC 1.14.21.6), C-5 desaturation; *ERG5* (EC 1.14.19.41), C-22 desaturation; *ERG4* (EC 1.3.1.71), delta-24 reduction; *SSR2* (EC 1.3.1.72), delta-24(25) reduction, *TtSTC1* (EC 4.2.1.123), *Tetrahymena thermophilia* squalene-tetrahymanol cyclase; *DWF5* (EC 1.3.1.21), delta-7 reduction; *DWF1/SSR1* (EC 1.3.1.72), delta-24(28) reduction; *SMT2* (EC 2.1.1.143), C-28 methylation; CYP710A1, C-22 desaturation. **b**, Gene edits required to create *Y. lipolytica* strains capable of producing non-native sterols that are tailored for honeybees.

To overcome this, we employed a sterol surrogate. The previously characterised *STC1* gene from *Tetrahymena thermophilia* encodes a squalene-tetrahymanol cyclase that converts squalene to the pentacyclic triterpenoid tetrahymanol (TET, Extended Data Fig. 1), which can substitute for ERGO function in yeasts^30–32^. TET is not expected to be detrimental to honeybees as pollen also contains diverse sterol intermediates and stanols that do not accumulate in honeybee tissues^20^. *TtSTC1* was introduced into the platform strain under the weak Pr*GPAT* promoter^33^ to drive minimal expression and limit diversion of carbon flux away from the sterol pathway. TET accounted for 14% of the sterol in the resulting TET strain (Extended Data Fig. 3a). By producing TET in *Y. lipolytica*, we were able to efficiently manipulate the sterol pathway. The subsequent elimination of ERGO production in our strains may also have helped to remove negative feedback inhibition on sterol biosynthesis.

We first engineered *Y. lipolytica* to produce the most important honeybee sterol, 24-MC. Redirecting the *Y. lipolytica* sterol pathway from ERGO to 24-MC required us to sequentially delete the endogenous genes *ERG4* and *ERG5* (sterol C-22 desaturase) and introduce a heterologous *DWF5*/*DHCR7* gene (delta-7 sterol reductase, Fig. 2). This strain (*erg4*Δ*erg5*Δ) was used to screen heterologous *DWF5*/*DHCR7* variants identified in the genomes of plants, animals, fungi, and bacteria. Ten diverse variants were selected and expressed under the strong Pr*TEFintron* promoter (Fig. 3a)^33^. The strain containing *Tetraselmis* (green algal phytoplankton) *DWF5* produced the highest content of 24-MC (42.1 mg/g DCW) and accumulated less TET and ergosta-5,7,24(28)-trienol intermediate than the next highest producing strain expressing *Candidatus* (bacterial pathogen of amoeba) *DWF5* (Fig. 3a).

**Fig. 3:**
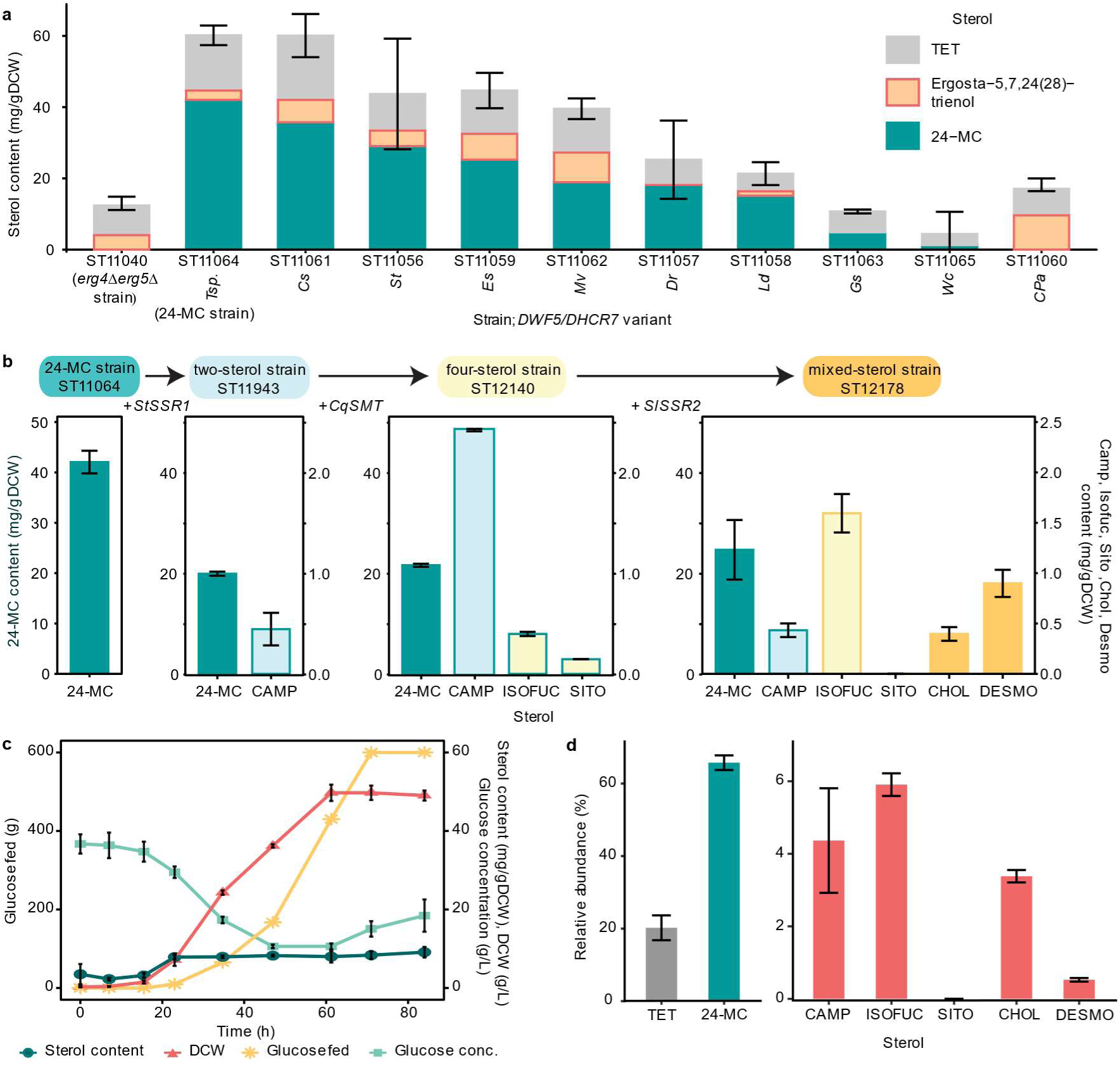
Engineering *Y. lipolytica* to produce key sterols required by honeybees. **a**, 24-MC production by engineered *Y. lipolytica* strains with different heterologous *DWF5/DHCR7* gene variants. Genes were selected from plants (*DWF5*), animals (*DHCR7*), algae, fungi, and bacteria, including *Solanum tuberosum*, *Danio rerio*, *Legionella drancourtii*, *Ectocarpus siliculosus*, *Candidatus* Protochlamydia amoebophila, *Coccomyxa subellipsoidea*, *Mortierella vertcillata*, *Glycine soja*, *Tetraselmis* sp. GSL018, and *Waddlia chondrophila*. Strains were compared for the content of 24-MC, TET and the main pathway intermediate ergosta- 5,7,24(28)-trienol (*n* = 3). Error bars represent ± s.d. for the total sterol content. **b**, Sterol composition of *Y. lipolytica* strains progressively engineered to produce a mixture of additional sterols required by honeybees (*n* = 3). TET and total sterol contents are shown in Extended Data Fig. 3. **c**, Time course of sterol content, DCW, glucose addition and broth glucose concentration during fed-batch fermentation of the mixed-sterol strain in 5 L bioreactors (*n* = 2). **d**, Sterol composition of the mixed-sterol strain after 84 h of fermentation (*n* = 2). Error bars represent ± s.d. Time courses of 5 L fermentation of the TET and W29 strains and sterol content from 5 L fermentation of the mixed-sterol, TET and W29 strains are shown in Extended Data Fig. 3.

The 24-MC strain was subsequently engineered to create a ‘mixed-sterol’ strain that produced small amounts of all the other sterols we observed in honeybees (Fig. 1b). We first introduced the previously characterised delta-24(28) sterol reductase (*SSR1*/*DWF1*) from potato *Solanum tuberosum*, which reduces the 24-MC side chain to produce CAMP (Fig. 2)^34^. By using a heterologous plant variant, we could regulate the expression level with the weak promoter Pr*DGA1*^33^ and obtain the C-24α/R epimer of CAMP found in plants, rather than the C-24β/S epimer as in ERGO (Extended Data Fig. 1). The resulting two-sterol strain produced 24-MC (19.9 mg/g DCW) and a small amount of CAMP (0.445 mg/g DCW, Fig. 3b). We then constructed a strain further capable of producing C-24 ethyl sterols, by introducing the C-28 sterol methyl transferase (*SMT*) from quinoa *Chenopodium quinoa* under the control of the Pr*GPAT* promoter. This four-sterol strain produced ISOFUC (0.396 mg/g DCW) and SITO (0.145 mg/g DCW) in addition to 24-MC (21.6 mg/g DCW) and CAMP (2.42 mg/g DCW, Fig. 3b).

To enable the production of CHOL and DESMO, the delta-24(25) sterol reductase (*SSR2*) from tomato *Solanum lycopersicum* was expressed under the Pr*GPAT* promoter in the four-sterol strain to generate the final mixed-sterol strain (Fig. 3b). Delta-24(25) sterol reductases are typically found in animals and reduce DESMO to CHOL but are also present in Solanaceous plants as part of the glycoalkaloid synthesis pathway^34^. When expressed in a 24-MC-producing *S. cerevisiae* strain, both DESMO and CHOL are produced^34^. This was similarly observed in our final mixed-sterol strain, which produced 24-MC (24.6 mg/g DCW), CAMP (0.433 mg/g DCW), ISOFUC (1.59 mg/g DCW), SITO (trace), CHOL (0.396 mg/g DCW) and DESMO (0.898 mg/g DCW) during small scale cultivation (Fig. 3b). The total detected sterol content decreased with each round of engineering from 60.2 mg/g DCW in the 24-MC strain to 28.0 mg/g DCW in the mixed-sterol strain (Extended Data Fig. 3b). The mixed-sterol strain was selected for cultivation in bioreactors.

We investigated the sterol and biomass production by cultivating the 24-MC strain in fed-batch mode in 250-mL controlled bioreactors (Ambr® 250). The 24-MC content reached 8.72-12.0 mg/g DCW, and the biomass reached 29.0-33.0 g DCW/L after 150 h of fermentation (Extended Data Fig. 3c, d). To produce biomass for testing in honeybee diets, we cultivated the mixed-sterol strain in fed-batch mode in 5 L bioreactors. After 84 h of fermentation the sterol content was 8.46-9.76 mg/g DCW, and the cell density reached 48.3-49.8 g DCW/L (Fig. 3c). The relative sterol content increased for the first 24 h and then remained stable. 24-MC comprised approximately 65% of the total sterol content (Fig. 3d). ISOFUC, CAMP, CHOL, DESMO and SITO together constituted 15% of the total sterols; TET contributed the remaining 20% (Fig. 3d). We also fermented the W29 strain and the TET strain in 5 L bioreactors for use as controls in feeding experiments, with the sterol contents reaching 0.0999-0.118 and 0.0991-0.204 mg/g DCW, respectively (Extended Data Fig. 3e, f).

Previous studies on non-native sterol production in yeast have achieved up to 36 mg/g DCW epi-campesterol in *Y. lipolytica* and 19.3 mg/g DCW epi-ergosterol in *S. cerevisiae* under fermentation conditions (Supplementary Table 1). We initially achieved 42.1 mg/g DCW of 24-MC during small scale cultivation, eight-fold more sterol per unit biomass than vegetable oils or floral pollen^20,35^, demonstrating the utility of the pre-engineered platform strain and the potential of *Y. lipolytica* as a host for sterol production. However, the sterol content declined with strain engineering and cultivation scale-up. Reduction in content has been observed in other studies with similar strains during bioreactor cultivation^36,37^. Nevertheless, pathway optimisation, capacity for sterol storage, and growth conditions for lipid accumulation can be significantly improved to increase the sterol content in *Y. lipolytica*^38^. Successful strategies in yeast have included regulation of cellular sterol homeostasis and enzyme re-localisation to lipid droplets^39,40^.

### Dietary sterols acquired from engineered yeast support extended brood production in honeybee colonies

The resulting yeast biomass from fermentation was dried and ground to create diets that could be tested in semi-field feeding trials with honeybee colonies. The diets were as follows: a ‘mixed-sterol strain diet’ (MxSt), and three control diets: a ‘TET strain diet’ (Tet), a ‘W29 strain diet’ (WT), and a ‘no yeast base diet’ (Base). Dried yeast biomass was incorporated at 20% w/w into an artificial diet formulation containing additional protein, fats, vitamins, and minerals required by honeybees. Because the mixed-sterol strain contained only trace SITO, all yeast diets were additionally supplemented with commercial phytosterol mix, containing approximately 12.0% w/w SITO (Supplementary Table 7). The Base diet was supplemented with additional commercial phytosterol mix to match the total sterol concentration in the MxSt diet (0.340% w/w).

We used a semi-field method with small honeybee colonies to measure the impact of the above diets upon brood production, in the absence of incoming pollen. We conducted feeding trials over three months and monitored diet consumption, sterol composition, colony size and brood status throughout (Fig 4, Extended Data Fig. 4-6). Every 15 days, we counted brood at the egg, larval and pupal (capped cell) stages in each colony, to assess the ability of each treatment to support brood production and development. In the first 45 days, an extreme heatwave resulted in a decline in bees of all ages across all colonies (Extended Data Fig. 7). Hence, we supplemented colonies with additional nurse bees on days 32 and 45. Nurse bees have endogenous sterols which can be used to feed larvae (Fig. 2)^18,21,41^. For this reason, we analysed the brood data starting from day 45 (t45) when we stopped adding nurses.

**Fig. 4:**
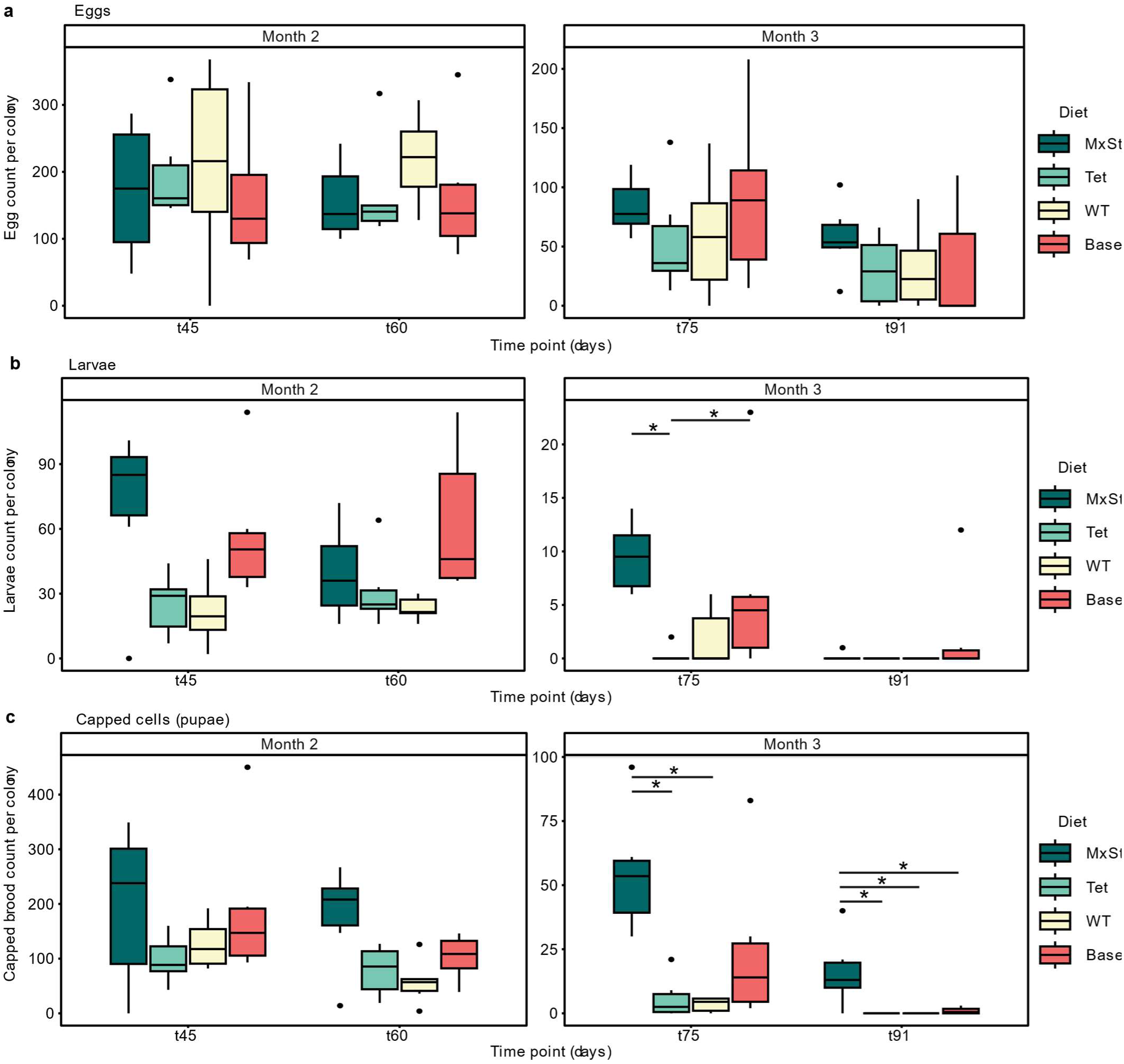
Semi-field trials to examine the impact of dietary sterols on honeybee colonies. Number of eggs (**a**), larvae (**b**) and capped brood cells (**c**) counted per colony for each treatment group for the last two months of the feeding trials. Eggs, larvae, and capped brood cells were counted in each colony at 15-day intervals (*n* = 6). Colonies were provided with a ‘mixed-sterol strain diet’ (MxSt) or one of three control diets: a ‘tetrahymanol strain diet’ (Tet), a ‘W29 strain diet’ (WT), and a ‘no yeast base diet’ (Base). Statistical analysis was performed by fitting 2- way negative binomial GLMMs followed by *post hoc* comparisons of estimated marginal means at each time point with Tukey adjustment (\**P* < 0.05, \*\**P* < 0.01, \*\*\**P* < 0.001). Brood counts from the first month of the feeding trials are shown in Extended Data Fig. 4. Note: the scales on months two and three differ, as brood production waned at the end of summer.

The number of eggs varied with time but not diet treatment group (two-way negative binomial GLMM, diet + time, time: χ^2^(3) = 48.3, *P* < 0.0001; diet: χ^2^(3) = 0.782, *P* = 0.854, Fig. 4a).

Feeding trials continued until mid-September when egg production in colonies begins to decline going into autumn. As such, there was a three-fold reduction in egg laying from t45-91 and a corresponding decrease in larvae, capped brood, and adults (Fig. 4, Extended Data Fig. 5). Overall, the number of larvae varied with diet treatment group, with the effect of diet becoming apparent by t75 (two-way negative binomial GLMM, diet*time, χ^2^(9) = 18.4, *P* = 0.0304, Fig. 4b). At t75, there were significantly more larvae in the MxSt diet treatment group than the Tet diet treatment group (*post hoc* comparison of estimated marginal means with Tukey adjustment, *P* = 0.0109). The Base diet treatment group also had more larvae than the Tet diet treatment group (*P* = 0.0295). By t91, larval counts were low in all colonies. The low brood production at this time point in the MxSt treatment was due to a seasonal effect that is also observed when colonies are fed with pollen (Extended Data Fig 8).

The diet significantly impacted the brood reared to pupation. As time passed, the impact of diet increased, such that colonies fed with the MxSt diet were more likely to rear larvae to the viable pupal stage (two-way negative binomial GLMM, diet*time, χ^2^(9) = 30.0, *P* = 0.0004,

Fig. 4c). By t75 there were significantly more pupae in the MxSt diet treatment group than the Tet (*P* = 0.0333) and WT (*P* = 0.0230) diet treatment groups. By the end of the experiment (t91), there were also significantly more pupae in the MxSt diet treatment group than the Base (*P* = 0.0328), Tet (*P* = 0.020) and WT (*P* = 0.020) diet treatment groups. There were still 10-40 capped brood cells in all but one colony in the MxSt diet treatment group, compared to 1-3 capped brood cells in three colonies in the Base diet treatment group and none in the control yeast diet groups. These data for diets without the required sterols are similar to our observations when colonies are starved of pollen: the rate of brood production is a function of the duration of nutrient starvation (Extended Data Fig 8). Eventually, in all colonies fed with diets lacking the required sterols (Base, Tet, and WT diets), the nurse bees did not have sufficient levels of key corporeal sterols necessary for brood rearing.

Although the number of pupae depended on the diet, sterol levels in the surviving pupae varied less than in the nurses and were not significantly different between treatment groups (Gaussian GLMM, diet*time, time: χ^2^(4) = 14.9, *P* = 0.005; diet: χ^2^(3) = 4.39, *P* = 0.222; Extended Data Fig. 6). The nurse bodies also showed less variation in sterol levels than the gut contents (Extended Data Fig. 6). However, neither the sterol content of the pupae nor the number of pupae correlated with the total sterol content of the nurse bodies (Extended Data Fig. 9). In the MxSt and Tet diet treatment groups, TET accumulated in the nurse guts (up to 42.4 μg/bee) but to a lesser extent in the nurse bodies (up to 18.7 μg/bee) and not at all in the pupae (Extended Data Fig. 6).

Pollen contains diverse sterols including cyclopropyl intermediates and stanols^20^, but our data indicate that only the delta-5 terminal intermediates of the sterol pathway tend to accumulate in honeybees. In yeast-feeding experiments with our mixed-sterol strain, we found that nurse workers transferred only the key sterols produced by the yeast (24-MC, ISOFUC, DESMO, CAMP, SITO, and CHOL) to the brood. Two diets contained the sterol surrogate TET, yet this was not found in the brood. This indicates that bees have selective mechanisms for enriching larval jelly with the key sterols. Our data validate the finding of Herbert *et al,* 1980, that honeybees likely require multiple sterols for continued brood production^10^. The most important sterols may be 24-MC and CHOL, as colonies fed sterol-free diets supplemented with 24-MC or CHOL produce more pupae than when supplemented with SITO or CAMP^42^.

We observed a clear benefit to the colonies with less than 0.5% w/w sterol inclusion in the diet. The yeast biomass is additionally a valuable source of protein, lipids, and vitamins^24,43^. Hence, it may be preferable to favour biomass over sterol yield when optimising fermentation to produce biomass for feed. We speculate that further engineering of lipid, protein, and terpenoid profiles to produce nutrients such as beneficial fatty acids, antioxidants, carotenoids, and fragrances^44–46^ could extend the suitability of *Y. lipolytica* biomass as a honeybee feed. *Y. lipolytica* biomass is regarded as safe for inclusion in food and feed, and engineered biomass has been established as feed in aquaculture^43,47,48^. Directly using inactivated yeast biomass in the diet circumvents the need for sterol extraction, which is expensive and inefficient, and minimises downstream processing. Such technology could also provide an avenue for developing feeds for other farmed arthropods.

## Methods

### Strains, culture conditions and chemicals

*Escherichia coli* strain DH5α was used for plasmid construction. *E. coli* was grown at 37 °C and 300 rpm in Lysogeny Broth (LB) liquid medium and at 37 °C on LB solid medium plates supplemented with 20 g/L agar. Ampicillin was supplemented at a concentration of 100 mg/L for plasmid selection.

The *Y. lipolytica* W29 strain (*MATa,* Y-63746 ARS Culture Collection, NCAUR, United States), and W29-derived platform strain ST9100 (*MATa ku70Δ::PrTEF1-cas9- TTef12::PrGPD-dsdA-TLip2 IntC_2-HMG1<-PrGPD-PrTefInt->ERG12 pCfB8823 IntC_3- SeACS<-PrGPD-PrTefInt->YlACL1 IntD_1-IDI1<-PrGPD-PrTefInt->ERG20,* melavonate- upregulated strain) were described in Arnesen *et al*, 2020^25^. The platform strain ST9100 was used to construct the sterol production strains. All strains are detailed in Supplementary Table 2.

*Y. lipolytica* was grown at 30 °C on yeast extract peptone dextrose (YPD) medium containing 10 g/L yeast extract, 20 g/L peptone, and 20 g/L glucose, supplemented with 20 g/L agar for preparation of solid media. For selection, either nourseothricin (250 mg/L) or hygromycin (400 mg/L) was added to the media. Cultivation of strains for sterol production was performed in YPD medium containing 80 g/L glucose. Chemicals were obtained, if not indicated otherwise, from Sigma-Aldrich/Merck (Germany). Nourseothricin was purchased from Jena BioScience GmbH (Germany).

### Plasmid construction

The coding sequences for delta-7 sterol reductase from *Solanum tuberosum*, (*StDWF5*, GenBank accession: BAQ55276.1), *Danio rerio* (*DrDHCR7*, accession: NP_958487.2), *Legionella drancourtii* (*LdDWF5*, accession: FJ197317.1), *Ectocarpus siliculosus* (*EsDWF5*, accession: CBN77313.1), *Candidatus* Protochlamydia amoebophila (*C*Pa*DWF5*, accession: KIC71363.1*), Coccomyxa subellipsoidea* (*CsDWF5*, accession: XM_005650286.1), *Mortierella vertcillata* (*MvDWF5*, accession: KFH65691.1), Glycine soja (*GsDWF5*, accession: XP_028244742.1); *Tetraselmis* sp. GSL018 (*T*sp*DWF5,* accession: JAC78771.1), and *Waddlia chondrophila* (*WcDHCR7*, accession: ADI39181.1), squalene-tetrahymanol cyclase from *Tetrahymena thermophilia* (*TtSTC1*, accession: XP_001026696.2), delta-24(25) sterol reductase from *Solanum lycopersicum* (*SlSSR2*, accession: BAQ55273.1), C-28 sterol methyl transferase from *Chenopodium quinoa* (*CqSMT*, accession: XP_021737090.1), and delta-24(28) sterol reductase from *Solanum tuberosum* (*StSSR1*, accession: AB839749.1) were codon-optimised for *Y. lipolytica* and synthesised as GeneArt Strings DNA fragments by Thermo Fisher Scientific. The codon-optimised sequences are listed in Supplementary Table 3.

The plasmids, BioBricks, and primers used in this study are listed in Supplementary Tables 4-6. BioBricks were amplified by polymerase chain reactions (PCR) using Phusion® U polymerase (Thermo Scientific). BioBricks were assembled into the EasyCloneYALI vectors with Uracil-Specific Excision Reagent (USER) cloning^33^. For marker-mediated gene deletion, upstream and downstream homology arms for relevant genes were synthesised as BioBricks by PCR amplification from platform strain ST9100 genomic DNA. Knock-out constructs were assembled from BioBricks by USER reaction as detailed in Supplementary Table 5. USER reactions were transformed into *E. coli* and correct assemblies were verified by Sanger sequencing (Eurofins).

### Yeast transformation

The yeast vectors were integrated into different previously characterised intergenic loci in the *Y. lipolytica* genome as described in Holkenbrink *et al*, 2018^33^. Integration vectors were digested with *Not*I enzyme (New England BioLabs) before lithium acetate transformation, as described by Holkenbrink *et al*, 2018^33^. Correct integration was verified by colony PCR using *Taq* DNA Polymerase Master Mix RED (Ampliqon) with vector-specific primers and primers complementary to the genomic region adjacent to the integration site^33^.

For marker-mediated gene deletion, transformants were selected on antibiotic supplemented plates and correct transformants confirmed by colony PCR. Marker removal was performed by transformation of the strains with a Cre-recombinase episomal vector^33^. Marker removal was confirmed by colony PCR.

### Yeast cultivation

Yeast strains were inoculated into 2.5 mL YPD in 24-deep well plates with air-penetrable lids (EnzyScreen, Netherlands). The plates were incubated at 30 °C with 300 rpm agitation for 24 hours (h). The optical density at 600 nm (OD600) was measured with an Implen P300 NanoPhotometer. The cultures were then diluted to OD600 0.1 in 2.5 mL fresh YPD-media with 80 g/L glucose and grown for a further 72 h at 30 °C with 300 rpm agitation. All cultivations were performed in triplicate. Cell dry weight was measured at the end of cultivation: 1 mL of culture broth was transferred into a pre-weighed 2 mL microcentrifuge tube, centrifuged (3000 g, 5 min) and the supernatant was discarded. The cells were washed twice with deionised water (1 mL). The cell pellet was dried at 60 °C for 7 days before the final weight was measured.

### Sterol analysis

For sterol extraction from yeast, 1 mL of culture broth was transferred into a 2 mL microcentrifuge tube, centrifuged and the supernatant was discarded. The cells were washed twice with deionised water (1 mL). The cell pellet was resuspended in 10% w/v methanolic potassium hydroxide (500 μL) and transferred to a 1 mL glass vial for saponification. The suspension was incubated at 70 °C for 2 h with vortexing at 15-minute intervals. The saponified samples were then vortexed and spiked with 50 μL of internal standard (1 mg/mL epicoprostanol in absolute ethanol). 500 μL of n-hexane was added to each sample for extraction of the free sterol component. Samples were vortexed and the organic phase transferred to a 2 mL microcentrifuge tube. The extraction step was repeated in a further 500 μL of n-hexane. The combined hexane phases were left overnight at room temperature for evaporation of the solvent. Sterol crystals remained in the tube.

For sterol analysis of the diets, each diet was sampled three times into pre-weighed 20 mL glass vials, and the weight of each sample was recorded. For sterol extraction from honeybee tissues, samples were first dried by freeze drying. Samples were dried at -48 °C under vacuum for four days. Dried samples were weighed and stored at -80 °C. For saponification, samples were first broken up with a spatula. For gut samples in 2 mL microcentrifuge tubes, samples were suspended in 500 μL 10% w/v methanolic potassium hydroxide. For honeybee tissue samples in 20 mL vials (pupae, nurse carcasses and queens), samples were suspended in 2.5 mL 10% w/v methanolic potassium hydroxide. Diet samples in 20 mL vials were suspended in 5 mL 10% w/v methanolic potassium hydroxide. Samples were incubated at 70 °C for 2 h in a water bath, with vortexing at 30-60 min intervals. The saponified samples were then spiked with 50 μL (gut samples) or 100 μL (diet, pupae, nurse carcasses and queen samples) of internal standard (1 mg/mL epicoprostanol in absolute ethanol). For extraction of the free sterol component, 500 μL (gut samples), 2.5 mL (pupae, nurse carcasses and queen samples) or 5 mL (diet samples) of n-hexane was added to each sample. Samples were vortexed and the organic phase transferred to a 2 mL (gut samples) or 7 mL (diet, pupae, nurse carcasses and queen samples) glass vial. The extraction step was repeated, and the combined hexane phases were left overnight at room temperature for evaporation of the solvent. The resulting extracts were resuspended in 500 μL (gut samples) or 1 mL (diet, pupae, nurse carcasses and queen samples) of n-hexane and vortexed. From each sample, a subsample of 250 μL was transferred to a 1.5 mL microcentrifuge tube and left at room temperature overnight for final drying.

Sterols were resuspended in 500 μL of pyridine, containing 20 μL *N,O*- Bis(Trimethylsilyl)acetamide (Merck) and incubated for four hours at 50 °C and then briefly vortexed before direct injection into an Agilent Technologies (Palo Alto, CA, USA) 8860 gas chromatograph connected to an Agilent Technologies 5977 MSD mass spectrometer (for gas chromatography–mass spectrometry (GC-MS)) and eluted over an Agilent HP-5MS column using a splitless injection at 250 °C with a standard GC program at 170 °C for 1 min, ramped to 280 °C at 20 °C min^-1^ and monitoring between 50 and 550 amu.

Sterols were identified by comparison of their retention time relative to cholesterol and mass spectra data available from the NIST (National Institute of Standards and Technology) mass spectral library and Zu *et al*, 2021^20^. Sterol identity in the final strain ST12178 was confirmed by comparison to authentic standards. Sterols were quantified by calculating the ratio of the peak area of the targeted sterol to that of the internal standard. The mass of each sterol in the sample was obtained by multiplying the ratio with the mass of the internal standard. Compound identification (using target ions) and quantification were carried out with ChemStation Enhanced Data Analysis (v.E.01.00).

### Bioreactor fed-batch cultivation

The Ambr® 250 system (Sartorius Stedim Biotech, Göttingen, Germany) was used to carry out 250 mL fed-batch fermentation in duplicate. The 24-MC strain ST11064 was re-streaked from glycerol stocks stored at -80 °C onto a YPD agar plate and incubated at 30 °C for 48 h. The preculture was prepared by inoculating strain ST11064 biomass from the plate into 50 mL YPD medium in a 250 mL shake flask and incubating at 30 °C for 24 h with 250 rpm agitation. Five mL of preculture was used to inoculate 115 mL batch media, to a starting OD600 of 0.25. For Ambr® 250 cultivation, the batch media comprised of mineral medium supplemented with yeast extract (10 g/L) and citric acid (20 g/L). The mineral medium was prepared with 0.5 g/L MgSO_4_⋅7H_2_O, 14.4 g/L KH_2_PO_4_, 0.1% (v/v) vitamin solution, 0.2% (v/v) trace metal solution as described in Jensen *et al*, 2014^49^, but with 3.4 g/L NH_4_Cl and glycerol as the carbon source (40 g/L).

Temperature was held constant at 30 °C. Dissolved oxygen was maintained above 20% by using a cascade of stirring speed ranging from 600 to 3000 rpm and aeration up to 1 vvm. The pH was maintained at 5.5 by the automatic addition of 1 M NaOH and 2.6 M H_3_PO_4_. Antifoam 204 (Sigma) was added automatically. Online measurements of acid and base addition, carbon dioxide evolution rate, dissolved oxygen and stirring speed were recorded for each reactor. The feed medium comprised 250 g/L glycerol. Feeding was automatically initiated once carbon dioxide evolution rate (CER) dropped below 50% at the end of the batch phase. Feeding was set to a constant rate of 0.9 mL/h. Samples were taken from each reactor every 6 h for the first 24 h, and then every 12 h, and immediately frozen until preparation for analysis. DCW and sterol content were determined from 1 mL samples as described above for small scale cultivation.

For larger scale fermentation, strains were cultivated by fed-batch fermentation in a 5 L bioreactor (BIOSTAT® B-DCU, Sartorius, Goettingen, Germany). All fermentations were carried out in duplicate. For each of strains W29, the TET strain ST11005 and the mixed-sterol strain ST12178, the strain was re-streaked from glycerol stocks onto a YPD agar plate and incubated at 30 °C for 24 h. The preculture was prepared by inoculating strain biomass from the plate into 50 mL YPD medium in a 250 mL shake flask and incubating at 30 °C for 24 h with 250 rpm agitation. The volume of pre-culture required to inoculate 2 L batch medium to a starting OD600 of 2.5 was centrifuged for 10 min at 4000 g and concentrated in 10 mL volume. This cell suspension was used to inoculate the bioreactors. The bioreactors were equipped with pH, pO2 and temperature probes. Temperature was held constant at 30 °C. Dissolved oxygen was maintained above 20% by adjusting stirring between 600 and 1200 rpm and aeration (via a horseshoe sparger) between 0.5 and 3 standard-litre per min (SPLM). The pH was kept at 5.5 by automatic addition of 5 M NaOH. Antifoam A (Sigma) was added as required.

The batch media comprised of mineral medium supplemented with yeast extract (20 g/L) and peptone (40 g/L). The mineral medium was prepared with 0.5 g/L MgSO_4_⋅7H_2_O, 14.4 g/L KH_2_PO_4_, 0.1% (v/v) vitamin solution, 0.2% (v/v) trace metal solution as described in Jensen *et al*, 2014^46^, 40 g/L glucose and 1 mL/L antifoam A (Sigma). The feed contained 5 g/L MgSO_4_⋅7H_2_O, 30 g/L KH_2_PO_4_, 1% (v/v) vitamin solution, 2% (v/v) trace metal solution as described in Sáez-Sáez *et al*, 2020^50^, with 300 g/L glucose. An exponential feeding profile was programmed, and feeding was initiated 24 h after inoculation. Feed rate, F (mL/h) followed the profile F=10*e^(0.05*t), where t = time (h) from the start of feeding. After 36 h of exponential feeding, the feed was switch to a constant rate of 75 mL/h until the end of fermentation.

Duplicate samples from each reactor were taken every 8 h for the first 24 h, and then every 12 h, to measure DCW, sterol content, OD600 and glucose concentration. DCW and sterol content were determined from 1 mL samples as described above for small scale cultivation. During fermentation, 1 mL culture broth was centrifuged, and the supernatant was used to measure glucose concentration via a glucose HK assay kit (Sigma). The supernatant was then filtered and frozen until further analysis. Glucose was later quantified on a Dionex Ultimate 3000 HPLC system equipped with a RI-101 Refractive Index Detector (Dionex) refractive index detector. An Aminex HPX-87H column 7.8 mm × 300 mm (Bio-Rad) with a Micro-Guard Cation H+ guard column 4.6 mm × 30 mm heated to 30 °C was injected with 10 µL sample. The mobile phase consisted of 5 mM H_2_SO_4_ with an isocratic flow rate of 0.6 mL/min, which was held for 15 min. HPLC data were processed using Chromeleon 7.2.9 software (Thermo Fisher Scientific). Glucose was identified and quantified using authentic standards. Glucose concentrations were calculated from peak area by extrapolation from a six-point calibration curve regression.

### Honeybee Diet Preparation

Yeast strains W29, the TET strain ST11005 and the mixed-sterol strain ST12178 were cultivated by 5-L fed-batch fermentation as described above. At the end of fermentation, the yeast biomass was recovered from the culture by centrifugation (4000 g, 20 min) and washed with deionised water. The biomass was heat-inactivated and dried (60 °C for a minimum of 24 h). The dried material was ground to a fine powder and stored at -20 °C until further use.

The yeast biomass cannot be subject to inactivation by autoclave or chemical treatment, as this will degrade the sterols present in the yeast. Incubation at 60 °C is commonly deemed sufficient for irreversible inactivation of the yeast and heat-inactivation of GMO yeast followed by feeding the inactivated yeast to live animals is a method that has been used previously in the UK^51^. Irreversible inactivation of the yeast was confirmed by a standard colony forming unit assay. The heat-inactivated dried yeast was dissolved at 10 mg/mL in water. The suspension was plated in serial dilution (100 μL plated of 10, 1, 0.1, 0.01 and 0.001 mg/mL suspensions) on YPD-agar and the plates were incubated at 30 °C for at least seven days. No growth of *Y. lipolytica* colonies was observed. The detection limit is one organism/mg material or 10^6^ viable organisms/kg material.

The yeast biomass was then incorporated into a meridic artificial diet at 20% w/w. Four diet types were prepared: a ‘mixed-sterol yeast’ diet containing the mixed-sterol strain ST12178, a ‘wild-type yeast’ diet containing strain W29, a ‘platform yeast’ diet containing the TET strain ST11005, and a base diet control without yeast supplementation. The base diet control was formulated to maintain total protein, sugar, sterol, and fat content at the same level as in the yeast supplemented diets. The content of this diet was a modified version of the diet of Sereia *et al.* 2013, Doi: 10.4025/actascianimsci.v35i2.16976. Specifically, the base diet contained 17% soy protein isolate (Soysol, MyVegan), 69.4% sugars (fructose, glucose, sucrose, and maltodextrin), 6% lipids, 6.50% deionised water, 0.100% vitamin and mineral supplement (Latshaw Apiaries), 0.6% commercial phytosterol mix (BulkSupplements; Supplementary Table 7), and 0.400% Carrageenan Kappa. The diets were divided into 50 g patties and stored at -20 °C until use. The yeast supplemented diets had the same proportion of protein (17%), carbohydrates (70%), and fats (6%) adjusted from the reagents of the base diet to accommodate nutrients present in the yeast. Specifically, the yeast supplemented diets contained the following: 20.0% dried yeast powder, 7.80% soy protein isolate (Soysol, MyVegan), 63.4% sugars, 0.1% commercial phytosterol mix (BulkSupplements, approximately 55% purity containing a mixture of sterols and stanols; Supplementary Table 9), 4% lipids, 4.20% deionised water, 0.100% vitamin and mineral supplement (Latshaw Apiaries), 0.400% Carrageenan Kappa (SpecialIngredients).

### Yeast feeding trials

For the sterol analysis of honeybee brood, used as a proxy for the natural sterol profile of honeybee castes, we sampled worker, drone, and queen pupae from naturally fed colonies in our apiary (John Krebs Field Station, Oxford). Worker pupae were directly collected from capped brood frames. Drone pupae were collected from capped drone comb (larger cell size). Queen pupae were reared by grafting young larvae (two to three days post-hatching) into Nicot Queen Rearing Cups (Paynes Bee Farms, Ltd.). These were placed in queen-less colonies in repurposed Styrofoam mini-nucleus hives (APIDEA®, Switzerland) for up to eight days until development to the capped brood stage. Tissues (three pupae per replicate, *n* = 5) were sampled into pre-weighed 20 mL glass vials and the fresh weight was recorded before storing at -80 °C until further analysis.

Feeding trials were conducted using honeybee colonies maintained in repurposed Styrofoam mini-nucleus hives made up of one brood box with five frames and a top feeder with a hole for patty delivery. Hives were maintained within a closed glasshouse environment designed to negate bee escape. Hives were distributed across two glasshouse rooms, with varying entrance orientation. Feeders with 30% w/v sugar solution and water were distributed inside the glasshouse and replenished as required. The in-hive and ambient temperature and humidity were recorded every 30 minutes using autonomous in-hive sensors (Supplementary Data 3). A misting system was installed to cool the temperatures in the glasshouse.

Initially, each hive contained 900-1200 adult bees, 2-3 frames brood/larvae/eggs and 1-2 frames of honey stores, but no bee bread. New mated queens were introduced in cages with beekeeping candy (Candito, PIDA, Italy), for slow release, three days prior to the start of the experiment.

Feeding trials were conducted over three months from June-September 2022 at the John Krebs Field Station, Oxford. Six hives were randomly assigned to each treatment group (*n* = 6). At the start of the experiment, diet patties were added via the top feeder and replaced throughout the experiment as required. Hive weight (after removal of diet patty) and patty weight were measured. The number of ‘bee seams’ (one seam defined as a continuous line of bees between adjacent frames, observed upon initial hive opening) and frames filled with honey (‘sugar stores’) were estimated by visual inspection. Presence of the mated queen, sugar stores, eggs, larvae, and capped brood were checked, and brood frames were photographed for subsequent counting. Eggs, larvae, and capped brood were counted using the Adobe Photoshop count tool. Full assessments of the hives were conducted every 15 days

Six days after each full assessment, hives were partially assessed with minimal disruption to the colony. Hive weight and patty weight were measured; bee seams and sugar stores were estimated by visual inspection. Presence of the mated queen, eggs, larvae, and capped brood were briefly checked. On days 21 and 45, hives with low populations (less than four bee seams) were topped up with orphanised nurse bees from mixed, naturally fed colonies.

At every assessment, nurse and pupae samples were taken from three hives from each treatment group. Six nurses and six pupae were collected from each of the sampled hives. Samples were collected into pre-weighed 20 mL glass scintillation vials. The fresh weights of the samples were measured, and the vials were stored at -80 °C. Nurse bees were dissected to separate the guts and gut contents from the rest of the tissues. This was done by partially thawing the samples on ice and pulling the guts from the abdomen by the stinger. The gut contents were transferred to a 2 mL microcentrifuge tube and the remaining tissues were returned to the 20 mL vial. Dissected samples were stored at -80 °C until further analysis.

### Pollen starvation trial

A semi-field pollen starvation trial was conducted from August-October 2023 at the John Krebs Field Station, Oxford. Colonies were housed in mini-nucleus hives set up identically to the yeast feeding trial and were maintained in one room of a mesh polytunnel purpose-built to prevent bee escape. One week before the start of the treatment, colonies were topped up with nurse bees from mixed, naturally fed colonies so that each box contained at least five bee seams, and re-queened where necessary. We used a mix of pre-existing colonies and newly established colonies, but all were fed pollen for at least one month prior to the start of the experiment and were producing brood.

Buckets of water and feeders with 30% w/v sugar solution were distributed inside the polytunnel. Colonies were supplied with pollen or candy patties. Pollen patties consisted of 80% multifloral pollen and 20% high-concentrated sugar syrup (∼70% w/v). Candy patties consisted of ∼80% beekeeping candy and ∼20% maltodextrin, which was added to slow consumption and reduce melting of the patty in the hive. Following a one month feeding period, colonies in treatment groups 1, 2, and 3 were deprived of pollen for the corresponding number of weeks and fed with candy only. The control group (0) was fed pollen throughout. Treatment groups were balanced across hive entrance orientations, colony strengths (bee seams), and position within the polytunnel.

We performed a partial assessment every week to measure patty weight and hive weight and estimate bee seams. Every two weeks, a full assessment recorded the presence of the mated queen, eggs, larvae, and capped cells, and the amount of sugar and pollen stores, and every frame was photographed. The photographs were used to count the number of cells with eggs, larvae, and pupae in each hive, using ImageJ (doi:10.1038/nmeth.2089).

### Statistical analysis

All graph plotting and statistical analysis was performed in R version 4.2.2^52^. Generalised linear models (GLMs) and generalised linear mixed models (GLMMs) were fitted using stats^52^ and glmmTMB^53^ packages. Models with non-significant interaction terms were re-run without the interaction term. Post hoc analysis was performed using the car^54^ and emmeans^55^ packages with Tukey adjustments for family-wise error rates.

The mean relative abundance of each of the major sterols in naturally fed honeybee pupae was calculated as a percentage of the total sterol. We compared sterol relative abundance across pupal castes using a GLM with quasi-binomial distribution (relative abundance ∼ sterol type*pupal caste). Sterol concentrations in pupae were calculated from the fresh weights of the pupal tissue. We compared sterol concentrations in pupal tissue across castes using GLMs with Gaussian distributions (sterol concentration ∼ sterol type*pupal caste). We compared the relative abundance of sterols in pollen using a GLM with quasi-binomial distribution (relative abundance ∼ sterol type). The coefficient of variation for each sterol was calculated by dividing the standard deviation by the mean relative abundance values of each sterol.

Counts for each brood type were compared across diet treatment groups using generalised linear mixed models, fitted to counts from t45 onwards, with “hive ID” as a random effect and negative binomial distributions (brood count ∼ diet*time + (1|hive ID)). For egg counts, the interaction term was dropped (egg count ∼ diet + time + (1|hive ID)).

Both the total diet provided to each colony and the total diet consumption by each colony were compared across diet treatment groups by fitting GLMs with Gaussian distribution (diet weight ∼ diet). Because hive weight correlated significantly with bee seams and consumption rate correlated significantly with hive weight for all treatment groups (Extended Data Fig. 8), the daily consumption rates within each interval were normalised by hive weight, as a proxy for colony size. The normalised consumption rates were compared across diet treatment groups using a GLMM, with “hive ID” as a random effect and Gaussian distributions (normalised consumption rate ∼ diet + time + (1|hive ID)).

The weight of each hive was compared across diet treatment groups using a GLMM, with “hive ID” as a random effect and Gaussian distributions (hive weight∼ diet + time + (1|hive ID)). The number of bee seams in each hive and the number of frames filled with honey were doubled to give integer values and compared across diet treatment groups using GLMMs, with “hive ID” as a random effect and Poisson distributions (2*bee seams ∼ diet + time + (1|hive ID); 2*sugar stores ∼ diet + time + (1|hive ID)).

For the sterol contents of nurses, nurse guts and pupae (μg per individual), GLMMs were fitted for each sample type within each sterol, with Gaussian distributions and “hive ID” as a random effect (sterol content ∼ diet*time + (1|hive ID)). Interaction terms were not significant in some models (total sterol in pupae, 24-MC in pupae, CAMP in nurses and nurse guts, ISOFUC in pupae, DESMO in pupae and nurse guts) and were dropped as appropriate.

We examined the relationships between variables measured during feeding trials by fitting GLMs (response variable ∼ predictor variable*diet). All models used Gaussian distributions, apart from when comparing capped brood counts to the total sterol content of nurse bodies from the same colony. In this case a negative binomial distribution was used. Where no significant interaction was found between diet treatment group and the predictor variable, the interaction term was dropped from the models (response variable ∼ predictor variable + diet). For each diet treatment group, linear regressions were fitted between the predictor and response variables of interest.

## Data Availability

All data will be made publicly available upon publication.

---

**Extended Data Fig. 1.**
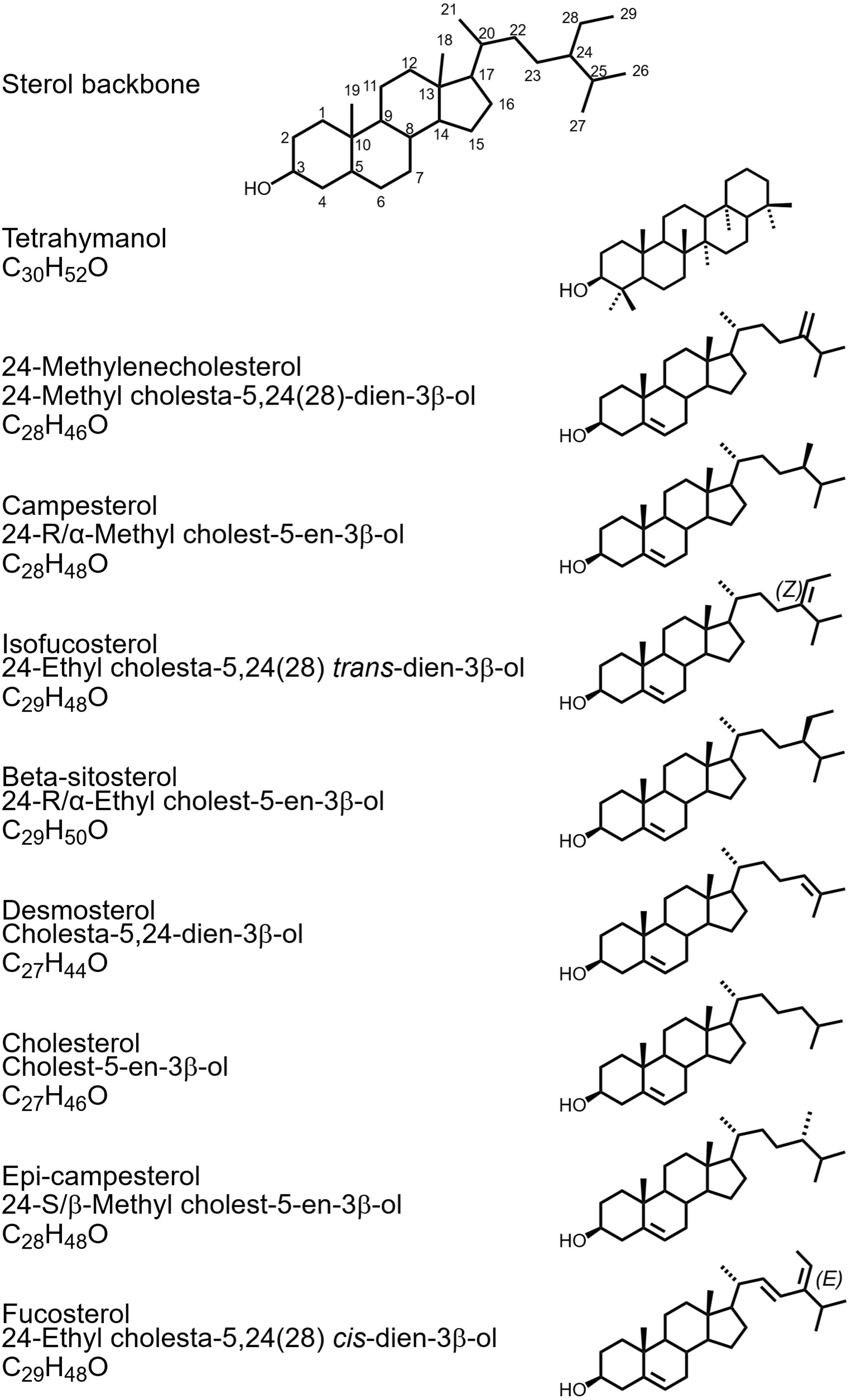
**Chemical structures, names, and formulae of sterols relevant to this study.**

**Extended Data Fig. 2.**
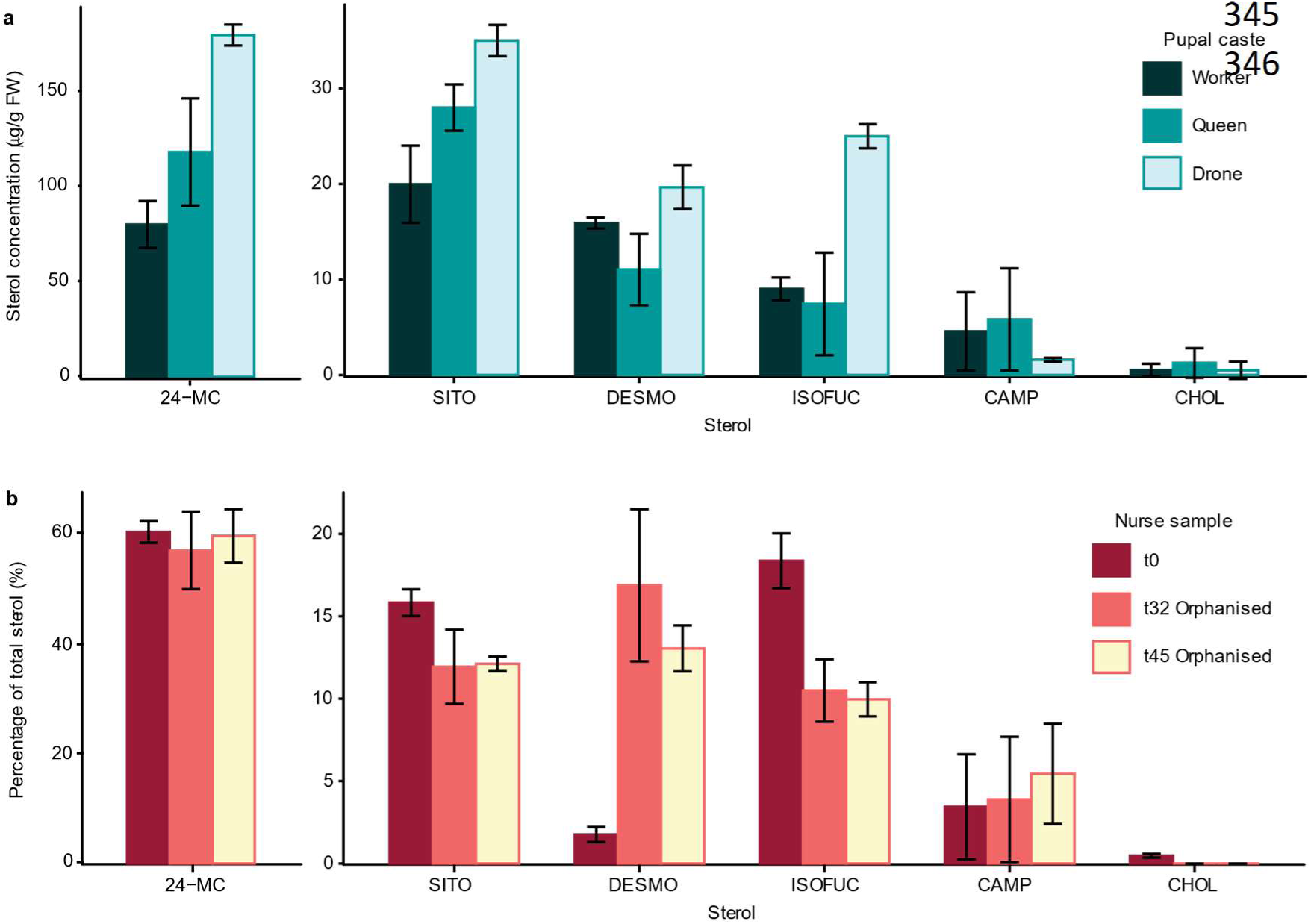
The major sterols in honeybee pupae from worker, queen, and drone bee castes and in nurse bees. **a**, Mean concentration of each of the major sterols per fresh weight of pupal tissue (*n* = 5). There was a significant interaction between sterol and pupal caste (Gaussian GLM, χ^2^(10) = 255, *P* < 0.0001). For each sterol, pairwise comparisons of estimated marginal means were carried out with Tukey *P*-value adjustment (\**P* < 0.05, \*\**P* < 0.01, \*\*\**P* < 0.001). 24-MC was more concentrated in drone than queen and worker pupal tissue. Isofucosterol (ISOFUC) was also more concentrated in drone than queen and worker pupal tissue. Sitosterol (SITO) was more concentrated in drone than worker pupae. **b**, Mean relative abundance of each of the major sterols in nurses from day 0, and orphanised top-up nurses from day 32 and day 45, given as a percentage of the total sterol (six pupae per sample, t0 nurses, *n* = 24; t32 orphanised nurses, *n* = 10; t45 orphanised nurses, *n* = 4). Error bars represent ± s.d.

**Extended Data Fig. 3.**
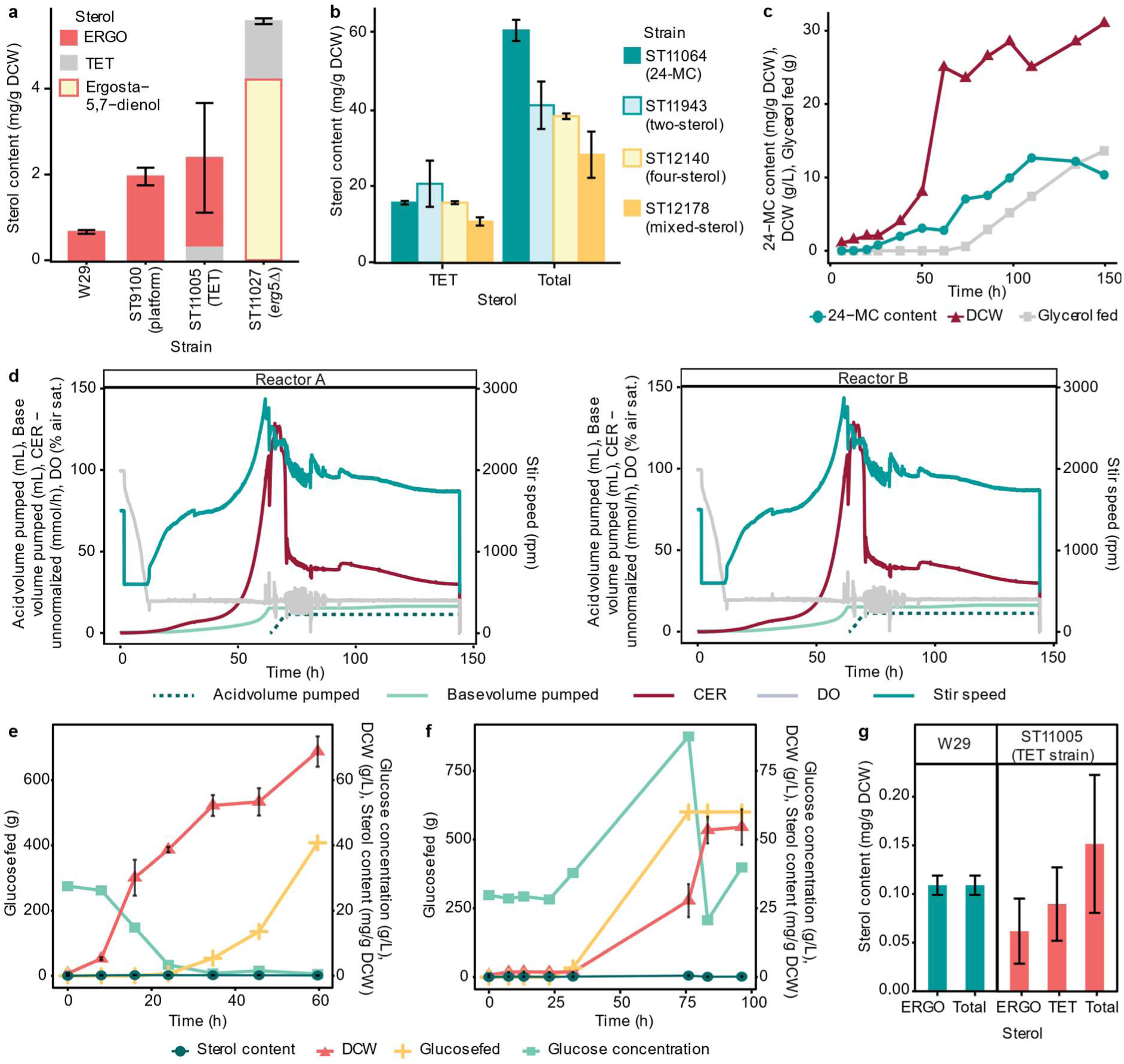
Characterisation and fermentation of engineered *Y. lipolytica* strains. **a**, Sterol content in W29, platform, TET and *erg5*Δ strains (*n* = 3). **b**, TET, and total sterol content in *Y. lipolytica* strains engineered to produce a mixture of non-native sterols (*n* = 3). **c**, Time course of sterol content, DCW and glycerol addition during fed-batch fermentation of the 24-MC strain in Ambr® 250 bioreactors (*n* = 2). **d**, Time course of operational parameters during fed-batch fermentation of the 24-MC strain in Ambr® 250 bioreactors. **e**, Time course of sterol content, DCW, glucose addition and glucose concentration in the broth during fed-batch fermentation of the W29 strain in 5-L bioreactors (*n* = 2). **f**, Time course of sterol content, DCW, glucose addition and glucose concentration in the broth during fed-batch fermentation of the TET strain in 5 L bioreactors (*n* = 2). **g**, Sterol composition of the W29 and TET strains after fed-batch fermentation in 5 L bioreactors (*n* = 2). Error bars represent ± s.d.

**Extended Data Fig. 4.**
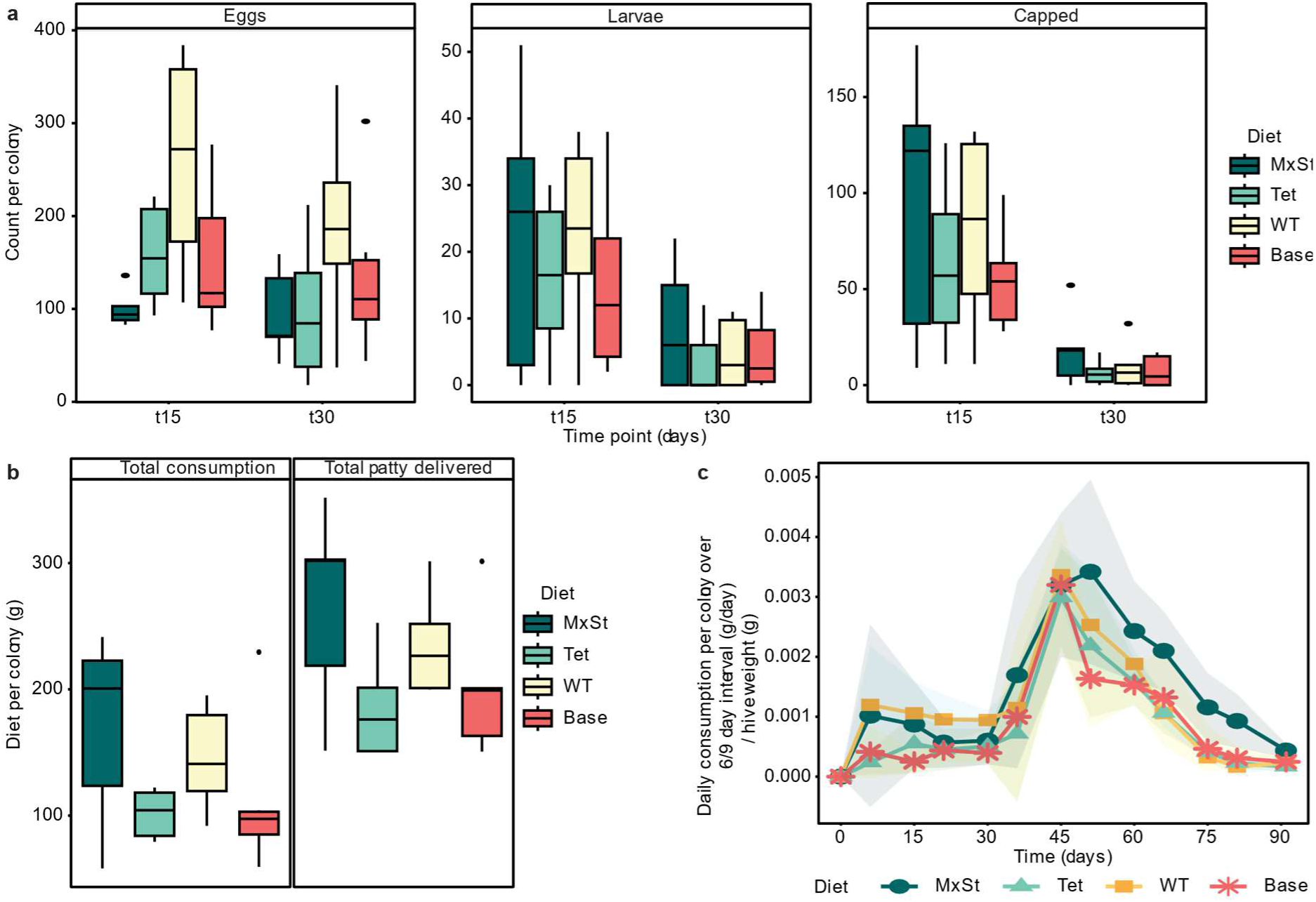
Initial colony brood status and diet consumption. **a**, Number of eggs, larvae and capped brood cells counted per hive for each treatment group for the first month of the feeding trials. Eggs, larvae, and capped brood cells were counted in each colony at 15-day intervals (*n* = 6). **b**, The total consumption (Gaussian GLM, χ^2^(3) = 3.15, *P* = 0.370) and total diet provided (χ^2^(3) = 5.74, *P* = 0.1252) to each colony over the course of the feeding trials did not vary significantly between treatment groups. **c**, Time course of diet consumption rate normalised by hive weight (*n* = 6). The normalised consumption rate varied with time (Gaussian GLMM, χ^2^(12) = 526, *P* < 0.0001) and diet treatment group (χ^2^(3) = 15.8, *P* = 0.0012). *Post hoc* pairwise comparison of estimated marginal means revealed that consumption rate was higher for colonies provided with the MxSt diet than those provided with the Tet (*P* = 0.0087) or Base (*P* < 0.0001) diets at t51. At t66, consumption rate for colonies provided with the MxSt diet was significantly greater than those provided with the Tet (*P* = 0.0388) or WT (*P* = 0.0385) diets. Error ribbons represent ± s.d.

**Extended Data Fig. 5.**
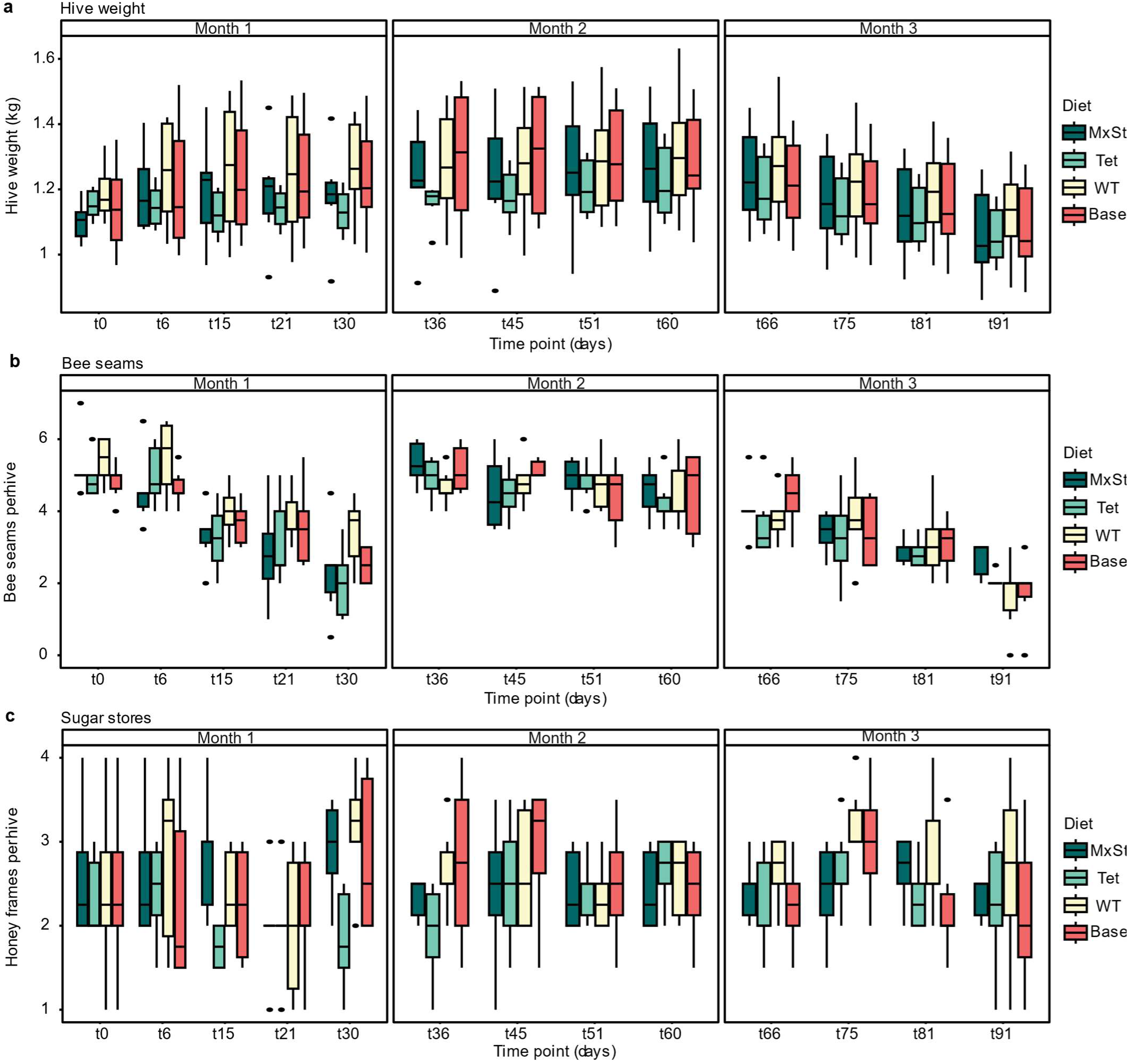
Feeding trial hive monitoring. **a**, Variation in hive weight during feeding trials (*n* = 6). Hive weight varied with time but not diet treatment group (Gaussian GLMM, time: χ^2^(12) = 176, *P* < 0.0001, diet: χ^2^(3) = 1.76, *P* = 0.623). **b**, Variation in number of bee seams during feeding trials (*n* = 6). Bee seams varied with time but not diet treatment group (Poisson GLMM (fitted to doubled count values), time: χ^2^(12) = 144, *P* < 0.0001; diet: χ^2^(3) = 1.38, *P* = 0.710). **c**, Variation in sugar stores during feeding trials (*n* = 6). Sugar stores did not vary significantly with time or diet treatment group (Poisson GLMM (fitted to doubled count values), time: χ^2^(12) = 8.67, *P* = 0.731; diet: χ^2^(3) = 3.11, *P* = 0.375).

**Extended Data Fig. 6.**
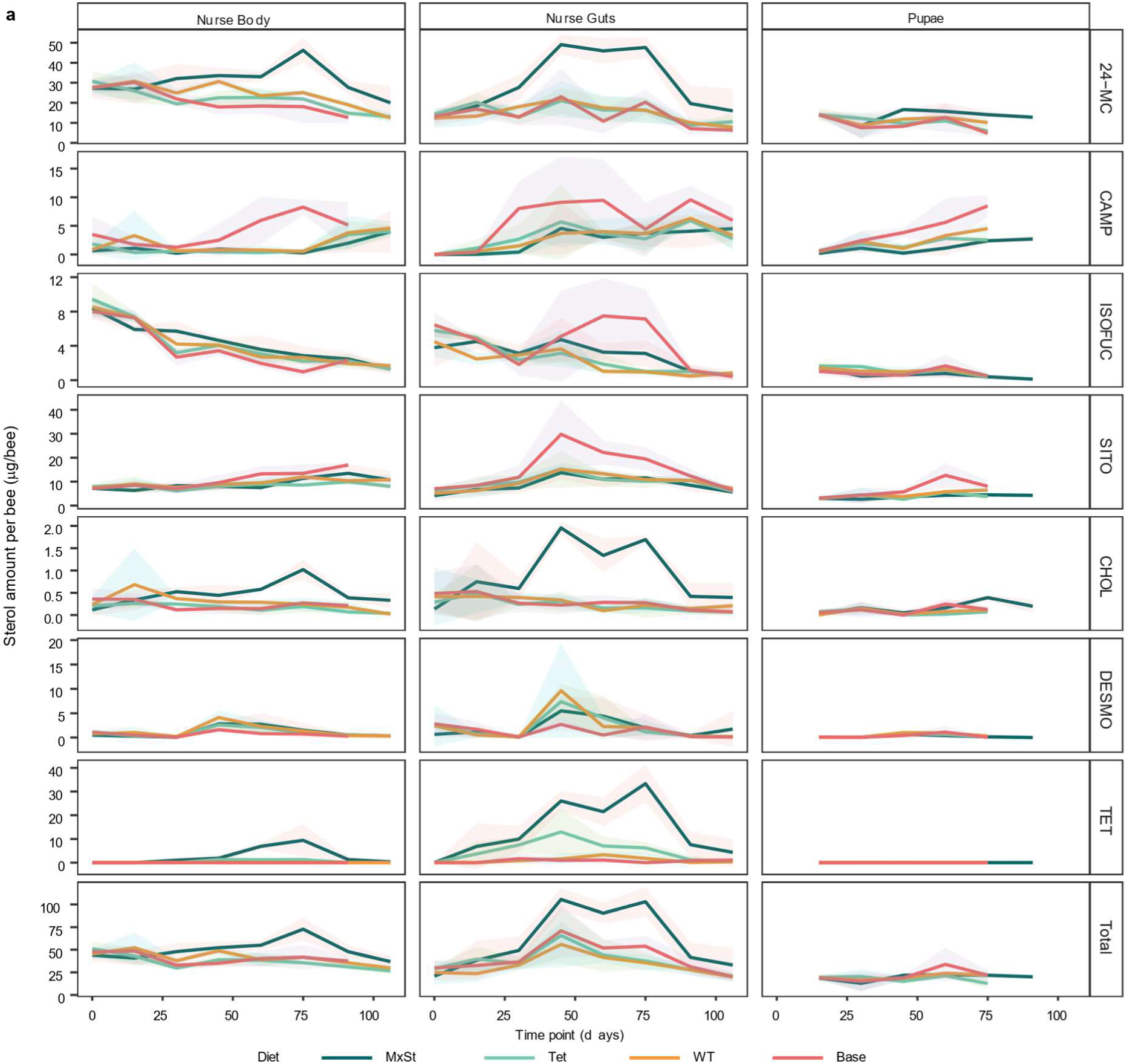
Time course of sterol content in nurse carcasses, nurse guts and pupae. **a**, Lines represent the average sterol content per bee (*n* = 3) for each treatment group (‘mixed-sterol strain diet’ (MxSt), ‘tetrahymanol strain diet’ (Tet), ‘W29 strain diet’ (WT), or ‘no yeast base diet’ (Base)). Error ribbons represent ± s.d. Full statistical analysis results are presented in Supplementary Data 2. From t45 to t75, there was significantly more sterol in the guts of nurses from the MxSt treatment group than those from any other treatment group (*post hoc* pairwise comparison of estimated marginal means, all *P* values < 0.0133). By t75 there was more sterol in the nurse bodies from the MxSt treatment group than all other treatment groups (all *P* values < 0.0001). Total sterol content in the pupae varied significantly with time but not diet treatment group (Gaussian GLMM, time: χ^2^(4) = 13.3, *P* = 0.0098; diet: χ^2^(3) = 3.50, *P* = 0.321). There was more TET in the nurse guts from the MxSt diet treatment group than all other treatment groups from t45-t60 (all *P* values <0.0005) and more TET in the nurse bodies from the MxSt diet treatment group than all other treatment groups at from t60-t75 (all *P* values <0.0001). Only trace amounts of TET were detected in the pupae.

**Extended Data Fig. 7.**
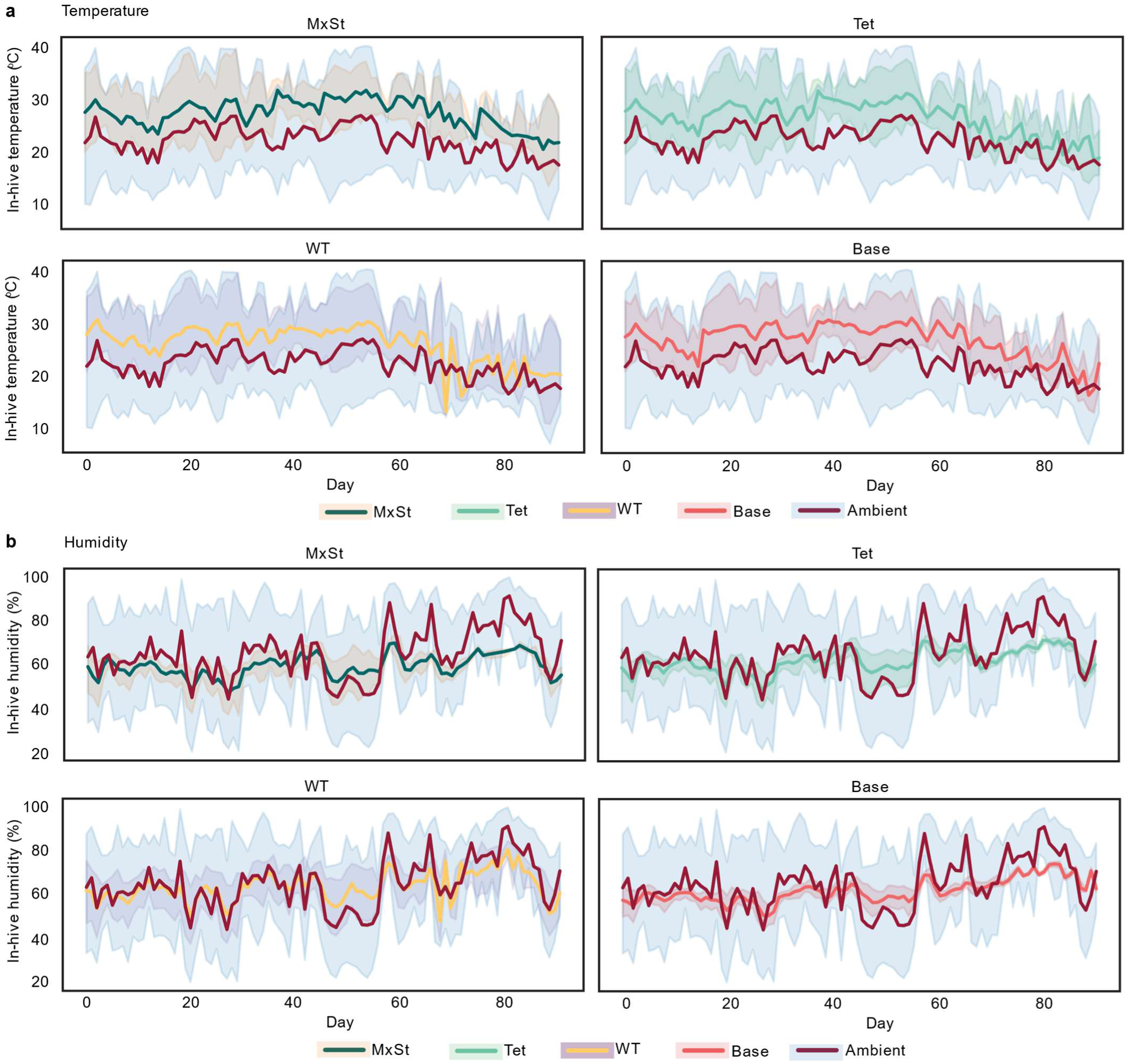
Temperature and humidity monitoring during feeding trials. Time course of temperature (**a**) and humidity (**b**) in mini-nucleus hives from each treatment group (‘mixed-sterol strain diet’ (MxSt), ‘tetrahymanol strain diet’ (Tet), ‘W29 strain diet’ (WT), or ‘no yeast base diet’ (Base)), and in the glasshouse (ambient). Lines represent the daily mean recordings. Ribbons depict the average maximum and minimum daily values. Note: extreme high temperatures occurred during days 26-29 and during days 49-55 which impacted colony brood production.

**Extended Data Fig. 8.**
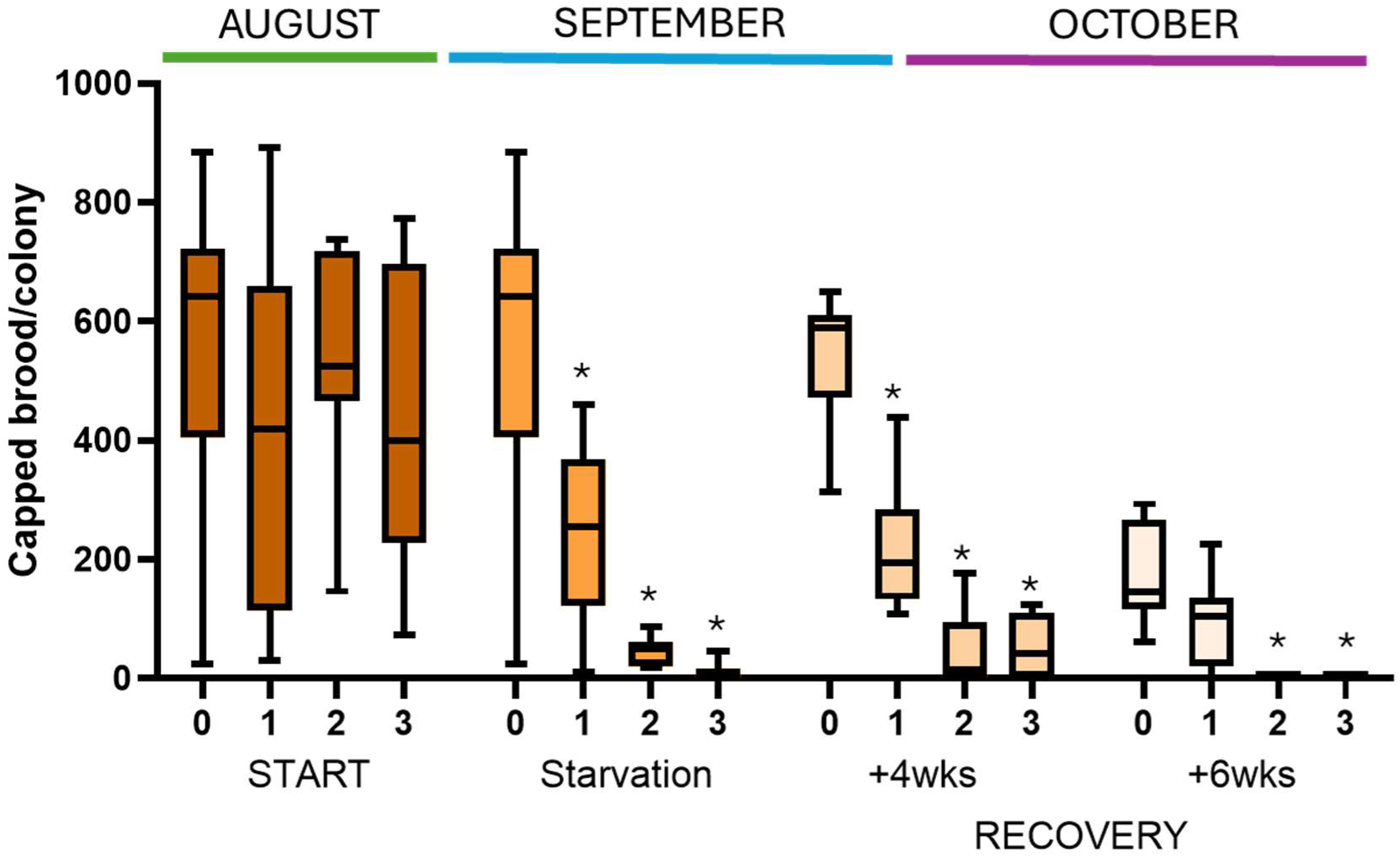
Pollen starvation combined with seasonal changes in a semi-field experiment leads to failed brood production. Brood production (count of capped cells/colony) was equal across all four treatments (0, 1, 2, 3, x-axis) at the start of the experiment (ANOVA, F_3, 26_ = 0.594, *P* = 0.625). Colonies starved for 1, 2 or 3 weeks (x-axis)) succumbed to seasonal reduction in brood production at a rate faster than the control (2-way ANOVA, time x treatment, F_6,77_ = 5.06, *P* < 0.001). The control (not starved, 0), like the other groups, exhibited a decline in brood production at the end of the season in October (Sidak, START vs +6wks for the control group, *P* = 0.003). The x-axis indicates the treatment (i.e. the number of weeks that the colonies were starved of pollen, 0, 1, 2, 3) over the three periods of the experiment (Start, Starvation period where no pollen was provided, and Recovery period of 4 weeks of pollen feeding or 6 weeks of pollen feeding. These data indicate that even when starved colonies are provided with pollen, it is not sufficient for them to recover from previous nutrient deprivation. * indicates a significant post-hoc comparison within time point of treatment vs control (*P* < 0.01). *n* = 8 colonies for the 1 and 3 week starvation treatments, *n* = 7 colonies for the pollen and 2 week starvation treatments.

**Extended Data Fig. 9.**
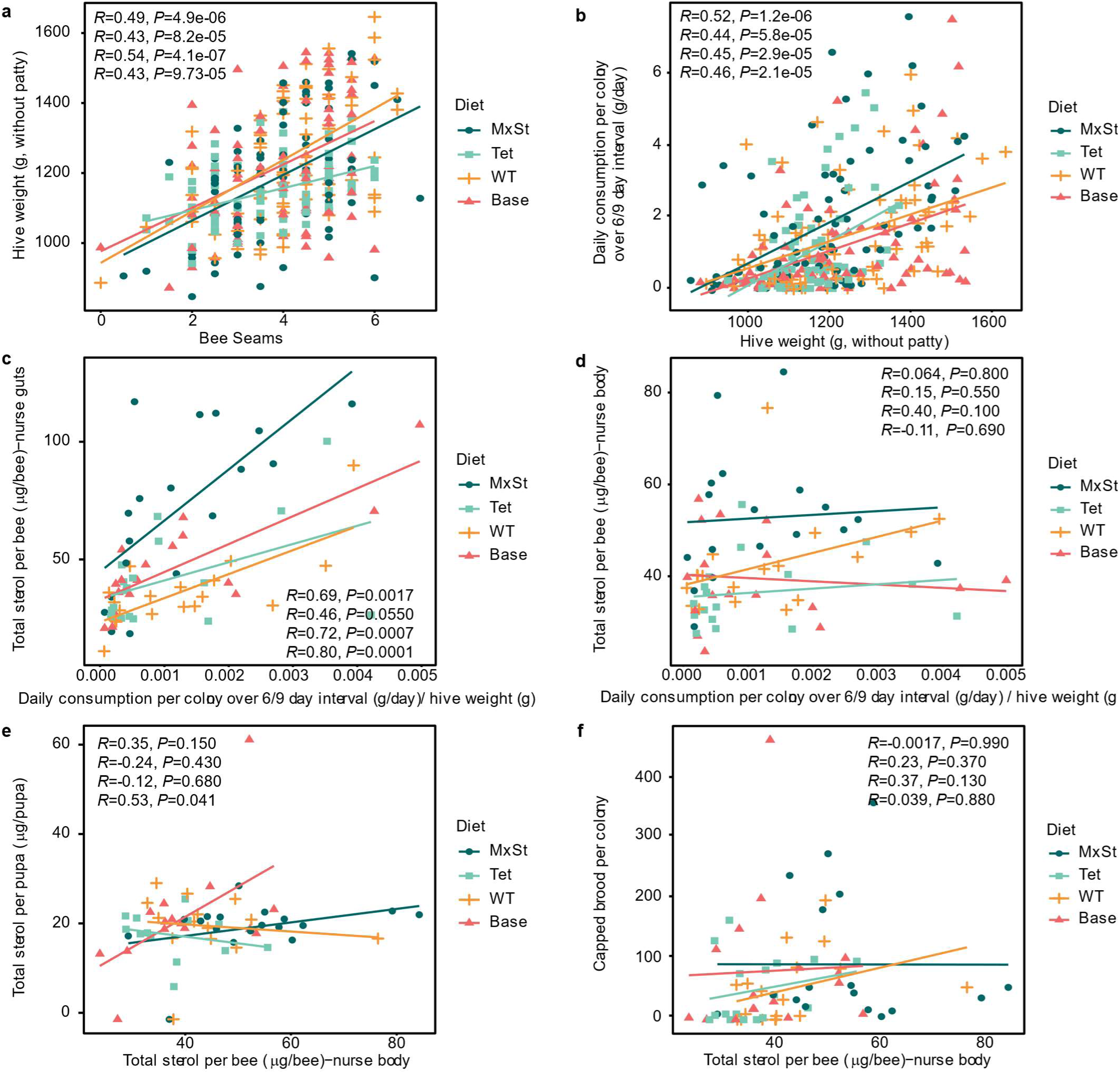
Regression analysis of feeding trial metrics. **a**, Hive weight varied significantly with bee seams (Gaussian GLM, χ^2^(1) = 83.9, *P* < 0.0001) and diet treatment group (χ^2^(3) = 12.9, *P* = 0.0049, ‘mixed-sterol strain diet’ (MxSt), ‘tetrahymanol strain diet’ (Tet), ‘W29 strain diet’ (WT), or ‘no yeast base diet’ (Base)). **b**, Consumption rate varied significantly with hive weight (Gaussian GLM, χ^2^(1) = 85.5, *P* < 0.0001) and diet treatment group (χ^2^(3) = 15.1, *P* = 0.0018). **c**, Total sterol per nurse gut varied significantly with normalised consumption rate (Gaussian GLM, χ^2^(1) = 43.6, *P* < 0.0001) and diet treatment group (χ^2^(3) = 36.2, *P* < 0.0001). **d**, Total sterol per nurse body did not vary significantly with normalised consumption rate (Gaussian GLM, χ^2^(1) = 0.844, *P* = 0.358) but did vary with diet treatment group (χ^2^(3) = 23.1, *P* < 0.0001). **e**, Sterol per pupa vs sterol per nurse body. There was a significant interaction between total sterol per nurse body and diet treatment group (Gaussian GLM, χ^2^(3) = 9.06, *P* = 0.0285). **f**, Capped brood cell counts did not vary with total sterol per nurse (negative binomial GLM, χ^2^(1) = 1.08, *P* = 0.300) or diet treatment group (χ^2^(3) = 1.83, *P* = 0.609). The Pearson correlation coefficient (*R*) and *P*-values are shown.

**Supplementary Table 1.** Yeasts engineered for non-native sterol production.

| Sterol | Host | Genotype<br>*giving highest<br>content only | Carbon source | Content/titre | Reference |
| --- | --- | --- | --- | --- | --- |
| Epi-campesterol<br>(‘ergosta-5-eneol’) | *S. cerevisiae*<br>FY1679-28C<br>(S288C<br>derivative) | *AtDHCR7* | Glucose/ethanol/galactose | 45% w/w total<br>sterol | 56 |
| Epi-campesterol<br>(‘ergosta-5-eneol’) | *S. cerevisiae*<br>FY1679-28C<br>(S288C<br>derivative) | *erg5Δ*<br>*AtDHCR7* | Glucose/ethanol/galactose | 0.3 mg/l/OD<br>(free sterol<br>only) | 57 |
| Epi-campesterol | *S. cerevisiae*<br>RH2881 | *erg5ΔDrDHCR7* | Glucose (2% w/v) | 98% w/w total<br>free sterol<br>Total sterol:<br>3.0 ug/10<sup>7</sup><br>cells | 58 |
| Epi-campesterol | *Y. lipolytica*<br>ATCC20124<br>9 | *erg5Δ*<br>*XlDHCR7* | Sunflower seed oil<br>(batch: 3.7% v/v, then<br>maintained < 2g/L) | 453 mg/L,<br>inferred<br>content: 20<br>mg/g DCW (5-L bioreactor,<br>fed-batch) | 59 |
| Campesterol | *S. cerevisiae*<br>BY4742 | *erg4Δerg5ΔArDWF1*<br>(plasmid) | Glucose (2% w/v)<br>Galactose (2% w/v) | Not quantified | 60 |
| Epi-campesterol | *Y. lipolytica*<br>ATCC20124<br>9 | *erg5ΔDrDHCR7*<br>↑*POX2* | Sunflower seed oil<br>(batch: 28.86 g/L, then<br>maintained < 2g/L) | 942 mg/L,<br>inferred<br>content: 36<br>mg/g DCW (5-L bioreactor,<br>fed-batch) | 61 |
| Epi-campesterol | *Y. lipolytica*<br>PO1f (ATCC<br>No. MYA-2613) | *erg5Δmfe1Δ*<br>*XlDHCR7* (2x) | Glucose (batch: 40 g/L,<br>maintained >10 g/L by<br>feeding 50% glucose<br>solution) | 837 mg/L,<br>inferred<br>content: 32<br>mg/g DCW (5-L bioreactor,<br>fed-batch) | 62 |
| Campesterol | *S. cerevisiae*<br>CENPK2.1D | *AtDWF7*<br>*AtDWF5*<br>*AtDWF1* ↑ (*ERG7*<br>*ERG8*<br>*ERG10*<br>*ERG12*<br>*ERG13*<br>*ERG19*<br>*ERG20*<br>*UPC2*<br>*tHMG1* (3x)) | Glucose (2% w/v) | 41 mg/L | 63 |
| Epi-campesterol | *S. cerevisiae*<br>YS5 | *erg5ΔXlDHCR7* | Glucose (20 g/L) | 178 mg/L (3<br>mL test tube) | 64 |
| Epi-campesterol | *S. cerevisiae*<br>BY-T30<br>(BY4742) | *StDWF5* ↑<br>(*ERG1*<br>*ERG8*<br>*ERG9*<br>*ERG10*<br>*ERG12*<br>*ERG13*<br>*ERG19*<br>*ERG20*<br>*IDII*<br>*tHMG1*) | Glucose (2% w/v) | 7.8<br>mg/L/OD600<br>(shake flask) | 65 |
| Cholesterol | *S. cerevisiae*<br>RH2881 | *erg5Δerg6Δ*<br>*DrDHCR7*<br>*DrDHCR24* | Glucose (2% w/v)<br>Molasses (0.75% v/v) | 96% w/w total<br>sterol,<br>1mg/g wet<br>weight of<br>yeast, 10 mg/L | 58 |
| Cholesterol | *P. pastoris*<br>CBS7435<br>*Δhis4Δku70* | *erg5Δerg6ΔDrDHCR7*<br>*DrDHCR24* | Glycerol (1% w/v)<br>Methanol (1% w/v) | 89% of total<br>sterol (shake<br>flask) | 66 |
| Cholesterol | *S. cerevisiae*<br>BY-T3<br>(BY4742) | *erg6Δatf2Δ*<br>*DrDHCR7*<br>*DrDHCR24*<br>↑<br>(*ERG1*<br>*ERG9*<br>*ERG20*) | Glucose (2% w/v)<br>Galactose (2% w/v) | 16 mg/L (250<br>mL shake<br>flask) | 67 |
| Cholesterol | *S. cerevisiae*<br>BY-T30<br>(BY4742) | *StDWF5 GgDHCR24*<br>↑<br>(*ERG1*<br>*ERG8*<br>*ERG9*<br>*ERG10*<br>*ERG12*<br>*ERG13*<br>*ERG19*<br>*ERG20*<br>*IDII*<br>*tHMG1*) | Glucose (2% w/v) | 8.5<br>mg/L/OD600<br>(shake flask) | 65 |
| Cholesterol<br>(intermediate in<br>diosgenin synthesis) | *S. cerevisiae*<br>(BY4742) | *atf2Δ*<br>*StDWF5 GgDHCR24*<br>*↓ERG6 VcCYP90B27*<br>*DzinCYP90G6*<br>*VcCYP94N1*<br>*VvCPR*<br>*VcCYP90B27*<br>*VcCYP94N1*<br>*VvCPR*<br>↑<br>(*ERG1*<br>*ERG8*<br>*ERG9*<br>*ERG10*<br>*ERG12*<br>*ERG13*<br>*ERG19*<br>*ERG20*<br>*IDII*<br>*tHMG1*) | Glucose (2% w/v)<br>(Fed to 5 g/L) | 170 mg/g<br>DCW (shake<br>flask),<br>5.5 g/L (5-L<br>fermentation) | 65 |
| Desmosterol | *S. cerevisiae*<br>RH2881 | *erg6ΔDrDHCR7* | Glucose (2% w/v) | 87% w/w total<br>free sterol<br>Total sterol:<br>3.4 ug/10<sup>7</sup><br>cells | 58 |
| Desmosterol | *S. cerevisiae*<br>BY4742 | *erg6ΔStDWF5* | Glucose (2% w/v)<br>Galactose (2% w/v) | Not quantified | 34 |
| β-Sitosterol | *S. cerevisiae*<br>CENPK2.1D | *erg4Δ*<br>*AtDWF7*<br>*AtDWF5*<br>*AtDWF1*<br>*SMT2 (plasmid)*<br>*are1Δ*<br>*are2Δ*<br>↑<br>(*ERG10*<br>*ERG12*<br>*ERG13*<br>*tHMG1*) |  | 2 mg/L | 63 |
| 7-Dehydrocholesterol | *S. cerevisiae*<br>RH2881 | *erg5Δerg6ΔDrDHCR2*<br>*4* | Glucose (2% w/v) | 86% w/w total<br>free sterol,<br>Total sterol:<br>3.2 ug/10<sup>7</sup><br>cells | 58 |
| 7-dehydrocholesterol | *S. cerevisiae*<br>RH2881 | *erg5Δerg6ΔGgDHCR2*<br>*4*<br>↑ (*ERG1*<br>*ERG2*<br>*ERG3*<br>*ERG11*<br>*Erg24*<br>*ERG25*<br>*ERG26*<br>*ERG27*) | Glucose (4% w/v)<br>Galactose (1% w/v) | 360.6 mg/L<br>(shake-flask) | 40 |
| 24-Epi-ergosterol | *S. cerevisiae*<br>CICC1746<br>(industrial<br>ergosterol<br>production<br>strain) | *ArDWF1*<br>*erg4Δ↑ (ERG5, YEH1,*<br>*YEH2, ARE2, UPC2)* | Glucose (batch: 10 g/L<br>glucose, feed: 500 g/L)<br>Ethanol | 2.76 g/L<br>19.27 mg/gDC<br>W<br>(2-L<br>bioreactor,<br>fed-batch) | 39 |
| 24-Methylenecholesterol | *S. cerevisiae*<br>BY4742 | *erg4Δerg5ΔStDWF5* | Glucose (2% w/v)<br>Galactose (2% w/v) | Not quantified | 34 |
| 24-Methylenecholesterol | *S. cerevisiae*<br>YS5 | *erg4Δerg5ΔXIDHCR7*<br>(2x) | Glucose (20 g/L) | 225 mg/L (250 mL shake flask) | 64 |
| 24-Methylenecholesterol | *Y. lipolytica*<br>ST9100<br>(W29 derivative) | *erg4Δ erg5Δ*<br>*SeACS TtSTC1*<br>*TspDWF7*<br>↑<br>(*HMG1*, *ERG12*, *ACLI*,<br>*ID11*, *ERG20*) | Glucose (8% w/v) | 42.2 mg/g DCW (24-deep well plate) | This study |
| Desmosterol + cholesterol | *S. cerevisiae*<br>BY4742 | *erg6ΔStDWF5*<br>*SISSR2* | Glucose (2% w/v)<br>Galactose (2% w/v) | Not quantified | 34 |
| 24-Methylenecholesterol + campesterol | *S. cerevisiae*<br>BY4742 | *erg4Δ*<br>*erg5ΔStDWF5*<br>*SISSR1* | Glucose (2% w/v)<br>Galactose (2% w/v) | Not quantified | 34 |
| 24-Methylenecholesterol + cholesterol + desmosterol (inferred) | *S. cerevisiae*<br>BY4742 | *erg4Δ*<br>*erg5ΔStDWF5*<br>*SISSR2* | Glucose (2% w/v)<br>Galactose (2% w/v) | Not quantified | 34 |
| 24-Methylenecholesterol + campesterol | *Y. lipolytica*<br>ST9100<br>(W29 derivative) | *erg4Δ erg5Δ*<br>*SeACS TtSTC1*<br>*TspDWF7*<br>*StSSR1*<br>↑<br>(*HMG1*, *ERG12*, *ACLI*,<br>*ID11*, *ERG20*) | Glucose (8% w/v) | 20.3 mg/g DCW (24-deep well plate) | This study |
| 24-Methylenecholesterol + campesterol + isofucoesterol + sitosterol | *Y. lipolytica*<br>ST9100<br>(W29 derivative) | *erg4Δ erg5Δ*<br>*SeACS TtSTC1*<br>*TspDWF7*<br>*StSSR1*<br>*CqSMT2*<br>↑<br>(*HMG1*, *ERG12*, *ACLI*,<br>*ID11*, *ERG20*) | Glucose (8% w/v) | 20.3 mg/g DCW (24-deep well plate) | This study |
| 24-Methylenecholesterol + campesterol + isofucoesterol + sitosterol + cholesterol + desmosterol | *Y. lipolytica*<br>ST9100<br>(W29 derivative) | *erg4Δ erg5Δ*<br>*SeACS TtSTC1*<br>*TspDWF7*<br>*StSSR1*<br>*CqSMT2*<br>*SIDHCR24*<br>↑<br>(*HMG1*, *ERG12*, *ACLI*,<br>*ID11*, *ERG20*) | Glucose (8% w/v)<br>(Bioreactor batch media<br>40 g/L glucose, feed<br>media 300 g/L glucose) | 24.6 mg/g DCW (24-deep well plate)<br>9.11 mg/g DCW (5 L bioreactor) | This study |

**Supplementary Table 2.** *Yarrowia lipolytica* strains used in this study.

| Strain name | Genotype | Parent strain/Reference | Elements used to construct strain |  |
| --- | --- | --- | --- | --- |
|  |  |  | gRNA vector | Integration vector/BioBrick |
| ST4842<br>'W29 strain' | MATa | *Y. lipolytica* W29 (MatA, ATCC<sup>®</sup>20460<sup>TM</sup>) strain Y-63746 from the ARS culture collection |  |  |
| ST6512 | MATa ku70Δ::PrTEF1->Cas9- TTef12::PrGPD->DsdA-TLip2 | ST4842, Marella *et al.* (2020)<sup>68</sup> |  |  |
| ST9100<br>'Platform strain' | MATa ku70Δ::PrTEF1->Cas9- TTef12::PrGPD->DsdA-TLip2 IntC_2-HMG1<-PrGPD-PrTefInt->ERG12 IntC_3-SeACS<-PrGPDPrTefInt->YIACL1 IntD_1-IDI1<-PrGPD-PrTefInt->ERG20 | ST6512, Arnesen *et al.* (2020)<sup>25</sup> |  |  |
| ST11005<br>'Tet strain' | MATa ku70Δ::PrTEF1-Cas9-TTef12::PrGPD-DsdA-TLip2 IntC_2-HMG1<-PrGPD-PrTefInt->ERG12 pCfB8823 IntC_3-SeACS<-PrGPD-PrTefInt->YIACL1 IntD_1-IDI1<-PrGPD-PrTefInt->ERG20 IntE_3-PrGPAT->TtSTC | ST9100 | pCfB6637 (pNat-YLgRNA3_IntE_3) | pCfB10494 (IntE3_PrGPAT->TtSTC) |
| ST11014 | MATa ku70Δ::PrTEF1-Cas9-TTef12::PrGPD-DsdA-TLip2 IntC_2-HMG1<-PrGPD-PrTefInt->ERG12 pCfB8823 IntC_3-SeACS<-PrGPD-PrTefInt->YIACL1 IntD_1-IDI1<-PrGPD-PrTefInt->ERG20 IntE_3-PrGPAT->TtSTC Δerg5 HphMX | ST11005 |  | BB5098 (Erg5_HphMX_K O) |
| ST11027 | MATa ku70Δ::PrTEF1-Cas9-TTef12::PrGPD-DsdA-TLip2 IntC_2-HMG1<-PrGPD-PrTefInt->ERG12 pCfB8823 IntC_3-SeACS<-PrGPD-PrTefInt->YIACL1 IntD_1-IDI1<-PrGPD-PrTefInt->ERG20 IntE_3-PrGPAT->TtSTC erg5Δ | ST11014 |  | pCfB6611 (pNat-PrExp-Cre) |
| ST11040 | MATa ku70Δ::PrTEF1-Cas9-TTef12::PrGPD-DsdA-TLip2 IntC_2-HMG1<-PrGPD-PrTefInt->ERG12 pCfB8823 IntC_3-SeACS<-PrGPD-PrTefInt->YIACL1 IntD_1-IDI1<-PrGPD-PrTefInt->ERG20 IntE_3-PrGPAT->TtSTC erg5Δ erg4Δ HphMX | ST11027 |  | BB5097 (Erg4_HphMX_K O) |
| ST11056 | MATa ku70Δ::PrTEF1-Cas9-TTef12::PrGPD-DsdA-TLip2 IntC_2-HMG1<-PrGPD-PrTefInt->ERG12 pCfB8823 IntC_3-SeACS<-PrGPD-PrTefInt->YIACL1 IntD_1-IDI1<-PrGPD-PrTefInt->ERG20 IntE_3-PrGPAT->TtSTC erg5Δ erg4Δ HphMX IntE_4-PrTEFInt->DHCR7_Stuberosum | ST11040 | pCfB6638 (pNat-YLgRNA2_IntE_4) | pCfB10249 (IntE4_PrTEFInt->DHCR7_Stuberosum) |
| ST11057 | MATa ku70Δ::PrTEF1-Cas9-TTef12::PrGPD-DsdA-TLip2 IntC_2-HMG1<-PrGPD-PrTefInt->ERG12 pCfB8823 IntC_3-SeACS<-PrGPD-PrTefInt->YIACL1 IntD_1-IDI1<-PrGPD-PrTefInt->ERG20 IntE_3-PrGPAT->TtSTC erg5Δ erg4Δ HphMX IntE_4-PrTEFInt->DHCR7_Drerio | ST11040 | pCfB6638 (pNat-YLgRNA2_IntE_4) | pCfB10250 (IntE4_PrTEFInt->DHCR7_Drerio) |
| ST11058 | MATa ku70Δ::PrTEF1-Cas9-TTef12::PrGPD-DsdA-TLip2 IntC_2-HMG1<-PrGPD-PrTefInt->ERG12 pCfB8823 IntC_3-SeACS<-PrGPD-PrTefInt->YIACL1 IntD_1-IDI1<-PrGPD-PrTefInt->ERG20 IntE_3-PrGPAT->TtSTC erg5Δ erg4Δ HphMX IntE_4-PrTEFInt->DHCR7_Ldrancou | ST11040 | pCfB6638 (pNat-YLgRNA2_IntE_4) | pCfB10251 (IntE4_PrTEFInt->DHCR7_Ldrancou) |
| ST11059 | MATa ku70Δ::PrTEF1-Cas9-TTef12::PrGPD-DsdA-TLip2 IntC_2-HMG1<-PrGPD-PrTefInt->ERG12 pCfB8823 IntC_3-SeACS<-PrGPD-PrTefInt->YIACL1 IntD_1-IDI1<-PrGPD-PrTefInt->ERG20 IntE_3-PrGPAT->TtSTC erg5Δ erg4Δ HphMX IntE_4-PrTEFInt->DHCR7_Esilicul | ST11040 | pCfB6638 (pNat-YLgRNA2_IntE_4) | pCfB10252 (IntE4_PrTEFInt->DHCR7_Esilicul) |
| ST11060 | MATa ku70Δ::PrTEF1-Cas9-TTef12::PrGPD-DsdA-TLip2 IntC_2-HMG1<-PrGPD-PrTefInt->ERG12 pCfB8823 IntC_3-SeACS<-PrGPD-PrTefInt->YIACL1 IntD_1-IDI1<-PrGPD-PrTefInt->ERG20 IntE_3-PrGPAT->TtSTC erg5Δ erg4Δ HphMX IntE_4-PrTEFInt->DHCR7_Cprotoch | ST11040 | pCfB6638 (pNat-YLgRNA2_IntE_4) | pCfB10253 (IntE4_PrTEF->DHCR7_Cprotoch) |
| ST11061 | MATa ku70Δ::PrTEF1-Cas9-TTef12::PrGPD-DsdA-TLip2<br>IntC_2-HMG1<-PrGPD-PrTefInt->ERG12 pCfB8823<br>IntC_3-SeACS<-PrGPD-PrTefInt->YIACL1 IntD_1-IDI1<-<br>PrGPD-PrTefInt->ERG20 IntE_3-PrGPAT->TtSTC erg5Δ<br>erg4Δ_HphMX IntE_4-PrTEFInt->DHCR7_Csubellip | ST11040 | pCfB6638 (pNat-YLgRNA2_IntE_<br>4) | pCfB10254<br>(IntE4_PrTEFInt-<br>>DHCR7_Csubell<br>ip) |
| ST11062 | MATa ku70Δ::PrTEF1-Cas9-TTef12::PrGPD-DsdA-TLip2<br>IntC_2-HMG1<-PrGPD-PrTefInt->ERG12 pCfB8823<br>IntC_3-SeACS<-PrGPD-PrTefInt->YIACL1 IntD_1-IDI1<-<br>PrGPD-PrTefInt->ERG20 IntE_3-PrGPAT->TtSTC erg5Δ<br>erg4Δ_HphMX IntE_4-PrTEFInt->DHCR7_Mverticillata | ST11040 | pCfB6638 (pNat-YLgRNA2_IntE_<br>4) | pCfB10255<br>(IntE4_PrTEFInt-<br>>DHCR7_Mverti<br>cillata) |
| ST11063 | MATa ku70Δ::PrTEF1-Cas9-TTef12::PrGPD-DsdA-TLip2<br>IntC_2-HMG1<-PrGPD-PrTefInt->ERG12 pCfB8823<br>IntC_3-SeACS<-PrGPD-PrTefInt->YIACL1 IntD_1-IDI1<-<br>PrGPD-PrTefInt->ERG20 IntE_3-PrGPAT->TtSTC erg5Δ<br>erg4Δ_HphMX IntE_4-PrTEFInt->DHCR7_Gsoja | ST11040 | pCfB6638 (pNat-YLgRNA2_IntE_<br>4) | pCfB10256<br>(IntE4_PrTEFInt-<br>>DHCR7_Gsoja) |
| ST11064<br>'24-MC<br>strain' | MATa ku70Δ::PrTEF1-Cas9-TTef12::PrGPD-DsdA-TLip2<br>IntC_2-HMG1<-PrGPD-PrTefInt->ERG12 pCfB8823<br>IntC_3-SeACS<-PrGPD-PrTefInt->YIACL1 IntD_1-IDI1<-<br>PrGPD-PrTefInt->ERG20 IntE_3-PrGPAT->TtSTC erg5Δ<br>erg4Δ_HphMX IntE_4-PrTEFInt->DHCR7_Tsp | ST11040 | pCfB6638 (pNat-YLgRNA2_IntE_<br>4) | pCfB10257<br>(IntE4_PrTEFInt-<br>>DHCR7_Tsp) |
| ST11065 | MATa ku70Δ::PrTEF1-Cas9-TTef12::PrGPD-DsdA-TLip2<br>IntC_2-HMG1<-PrGPD-PrTefInt->ERG12 pCfB8823<br>IntC_3-SeACS<-PrGPD-PrTefInt->YIACL1 IntD_1-IDI1<-<br>PrGPD-PrTefInt->ERG20 IntE_3-PrGPAT->TtSTC erg5Δ<br>erg4Δ_HphMX IntE_4-PrTEFInt->DHCR7_Wchondrophi | ST11040 | pCfB6638 (pNat-YLgRNA2_IntE_<br>4) | pCfB10258<br>(IntE4_PrTEFInt-<br>>DHCR7_Wchon<br>drophi) |
| ST11943<br>'Two-sterol<br>strain' | MATa ku70Δ::PrTEF1-Cas9-TTef12::PrGPD-DsdA-TLip2<br>IntC_2-HMG1<-PrGPD-PrTefInt->ERG12 pCfB8823<br>IntC_3-SeACS<-PrGPD-PrTefInt->YIACL1 IntD_1-IDI1<-<br>PrGPD-PrTefInt->ERG20 IntE_3-PrGPAT->TtSTC erg5Δ<br>erg4Δ_HphMX IntE_4-PrTEFInt->DHCR7_Tsp<br>IntF3_PrDGA1->StSSR1 | ST11064 | pBP8003 (pNat-YLgRNA4-IntF_3) | pCfB10948<br>(IntF3_PrDGA1-<br>>StSSR1) |
| ST12140<br>'Four-sterol<br>strain' | MATa ku70Δ::PrTEF1-Cas9-TTef12::PrGPD-DsdA-TLip2<br>IntC_2-HMG1<-PrGPD-PrTefInt->ERG12 pCfB8823<br>IntC_3-SeACS<-PrGPD-PrTefInt->YIACL1 IntD_1-IDI1<-<br>PrGPD-PrTefInt->ERG20 IntE_3-PrGPAT->TtSTC erg5Δ<br>erg4Δ_HphMX IntE_4-PrTEFInt->DHCR7_Tsp<br>IntF3_PrDGA1->StSSR1 IntE1_PrGPAT->SMT2_Cq | ST11943 | pCfB6633 (pNat-YLgRNA2_IntE_<br>1) | pCfB10851<br>(IntE1_PrGPAT-<br>>SMT2_Cquinoa) |
| ST12178<br>'Mixed-sterol<br>strain' | MATa ku70Δ::PrTEF1-Cas9-TTef12::PrGPD-DsdA-TLip2<br>IntC_2-HMG1<-PrGPD-PrTefInt->ERG12 pCfB8823<br>IntC_3-SeACS<-PrGPD-PrTefInt->YIACL1 IntD_1-IDI1<-<br>PrGPD-PrTefInt->ERG20 IntE_3-PrGPAT->TtSTC erg5Δ<br>erg4Δ_HphMX IntE_4-PrTEFInt->DHCR7_Tsp<br>IntF3_PrDGA1->StSSR1 IntE1_PrGPAT->SMT2_Cq<br>IntE5_PrGPAT->DHCR24_Sl | ST12140 | pCfB10783<br>(pNatYlgRNA-IntE_5) | pCfB10853<br>(IntE5_PrGPAT-<br>>DHCR24_Slyco<br>persic) |

**Supplementary Table 3.**
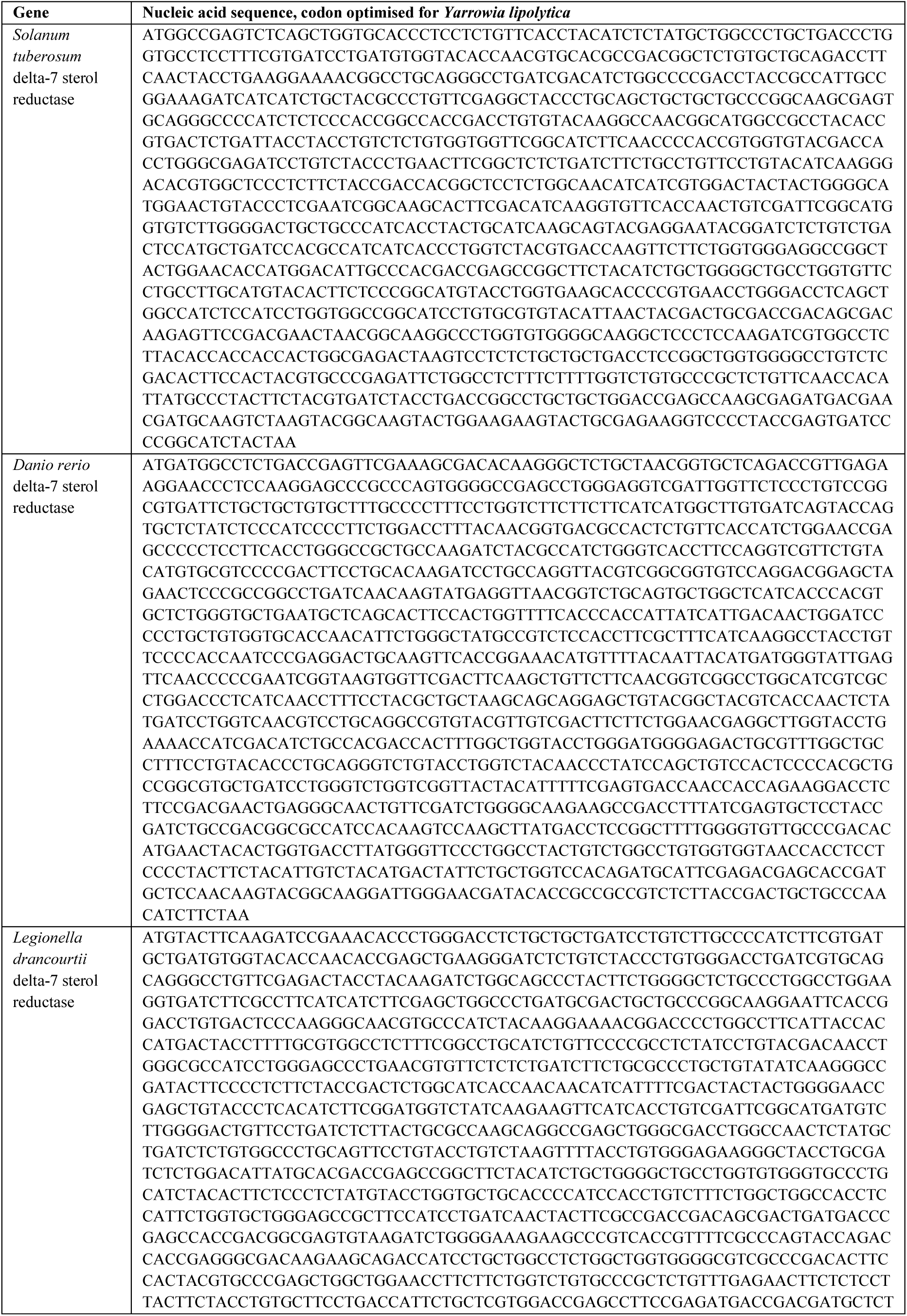
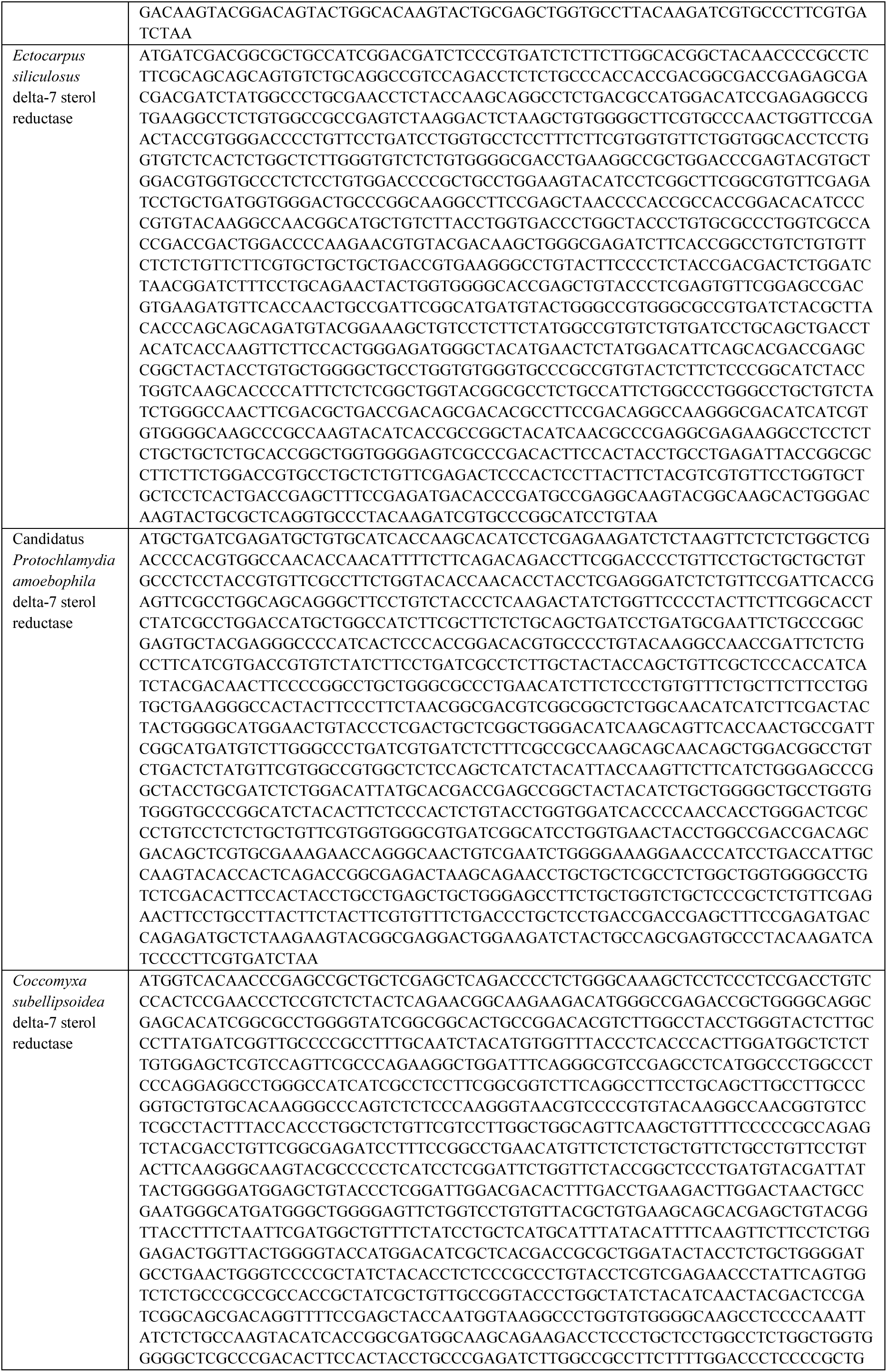
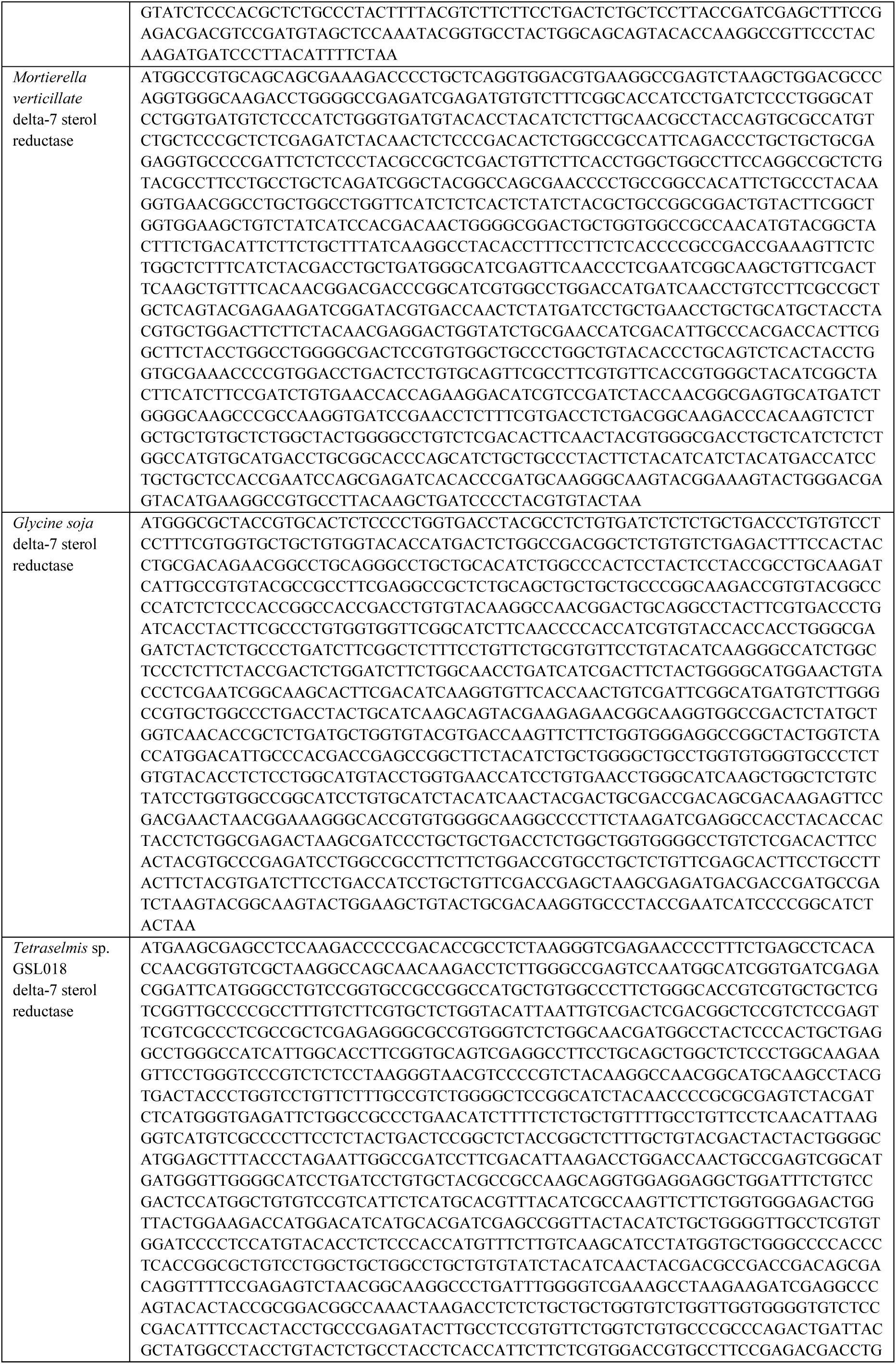
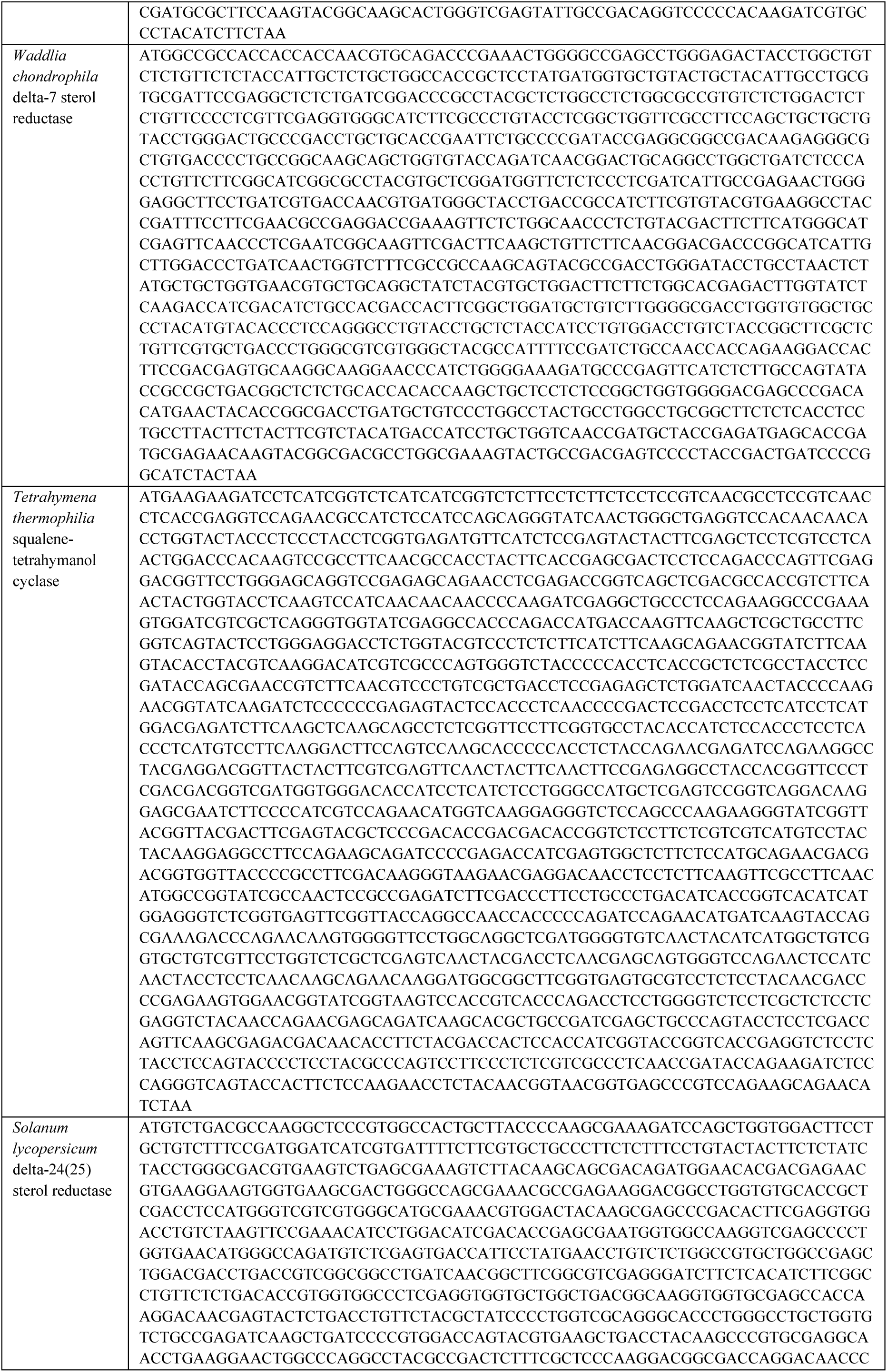
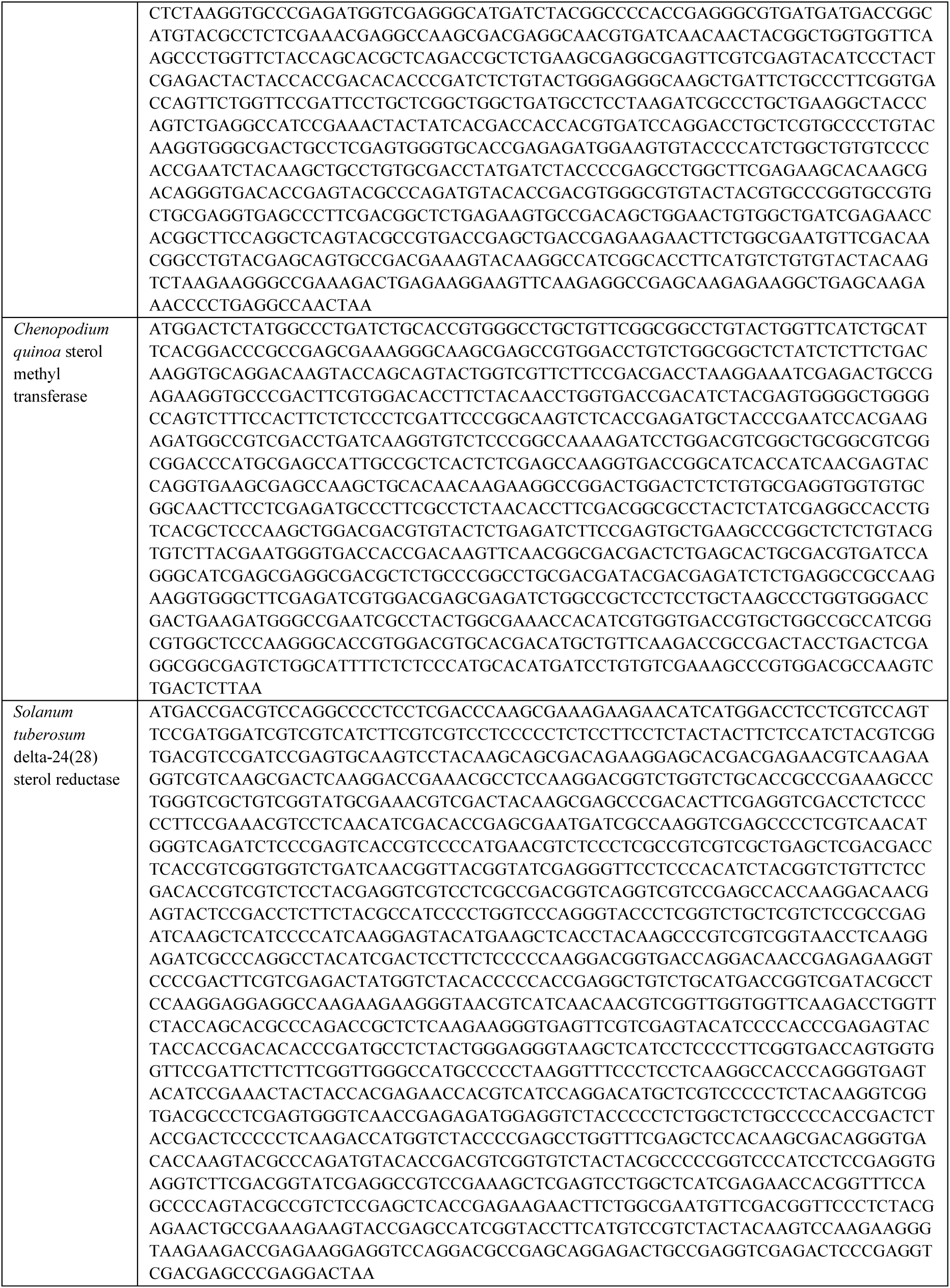
Codon-optimised sequences of heterologous genes used in this study.

**Supplementary Table 4.**
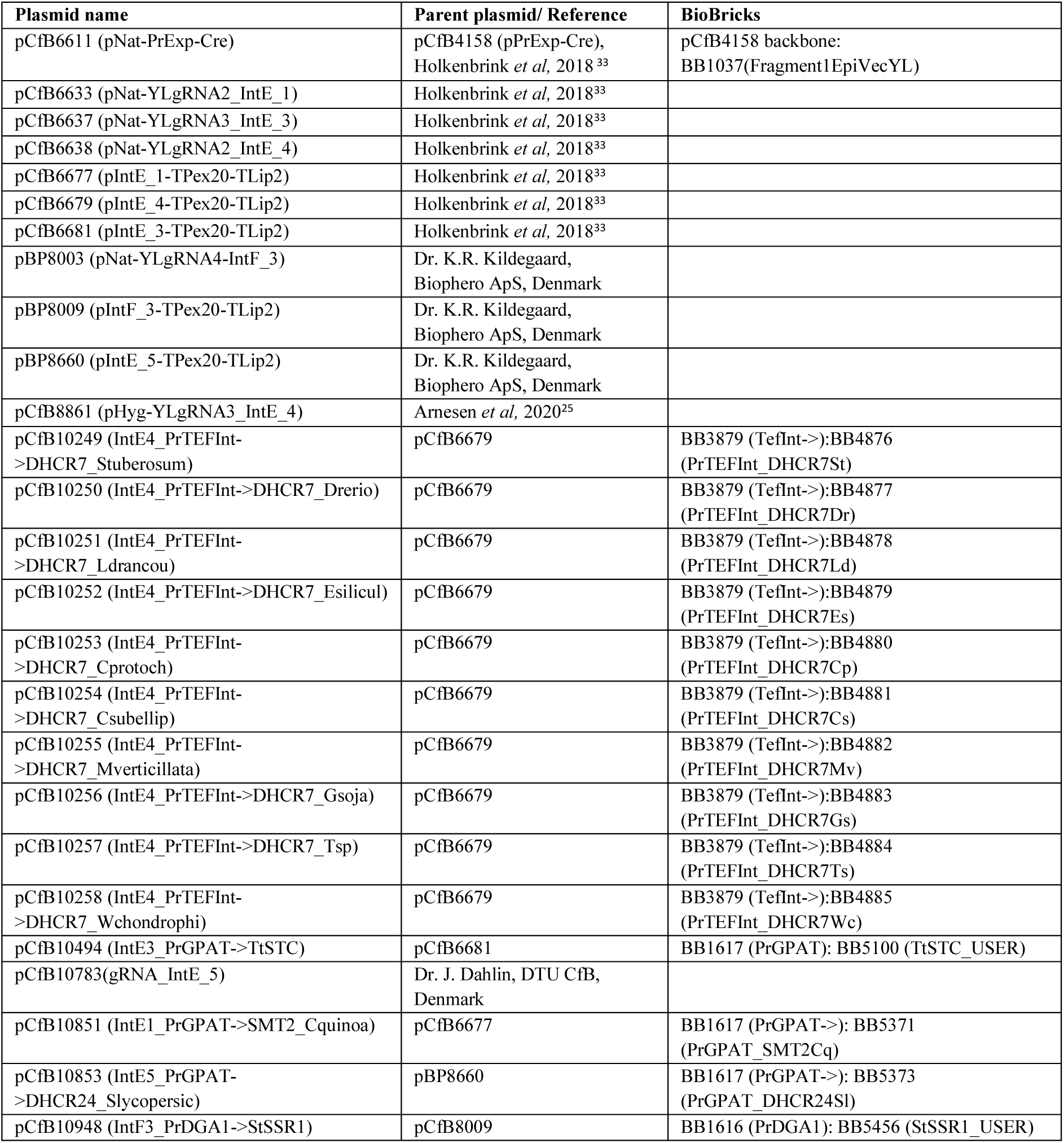
Plasmids used in this study.

**Supplementary Table 5.**
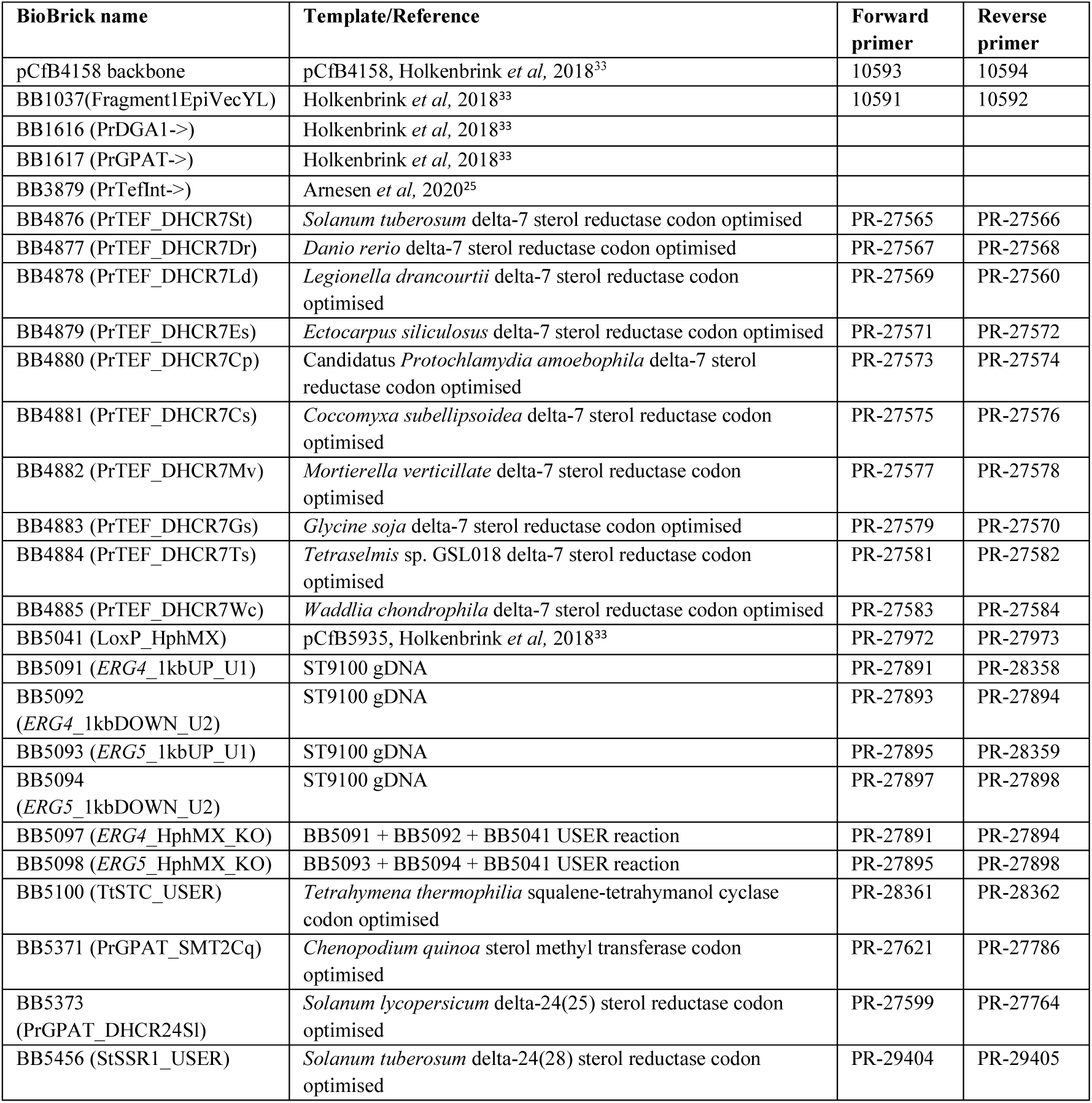
BioBricks used in this study.

| BioBrick name | Template/Reference | Forward primer | Reverse primer |
| --- | --- | --- | --- |
| pCfB4158 backbone | pCfB4158, Holkenbrink *et al*, 2018<sup>33</sup> | 10593 | 10594 |
| BB1037(Fragment1EpiVecYL) | Holkenbrink *et al*, 2018<sup>33</sup> | 10591 | 10592 |
| BB1616 (PrDGA1->) | Holkenbrink *et al*, 2018<sup>33</sup> |  |  |
| BB1617 (PrGPAT->) | Holkenbrink *et al*, 2018<sup>33</sup> |  |  |
| BB3879 (PrTefInt->) | Arnesen *et al*, 2020<sup>25</sup> |  |  |
| BB4876 (PrTEF_DHCR7St) | *Solanum tuberosum* delta-7 sterol reductase codon optimised | PR-27565 | PR-27566 |
| BB4877 (PrTEF_DHCR7Dr) | *Danio rerio* delta-7 sterol reductase codon optimised | PR-27567 | PR-27568 |
| BB4878 (PrTEF_DHCR7Ld) | *Legionella drancourtii* delta-7 sterol reductase codon optimised | PR-27569 | PR-27560 |
| BB4879 (PrTEF_DHCR7Es) | *Ectocarpus siliculosus* delta-7 sterol reductase codon optimised | PR-27571 | PR-27572 |
| BB4880 (PrTEF_DHCR7Cp) | Candidatus *Protochlamydia amoebophila* delta-7 sterol reductase codon optimised | PR-27573 | PR-27574 |
| BB4881 (PrTEF_DHCR7Cs) | *Coccomyxa subellipsoidea* delta-7 sterol reductase codon optimised | PR-27575 | PR-27576 |
| BB4882 (PrTEF_DHCR7Mv) | *Mortierella verticillate* delta-7 sterol reductase codon optimised | PR-27577 | PR-27578 |
| BB4883 (PrTEF_DHCR7Gs) | *Glycine soja* delta-7 sterol reductase codon optimised | PR-27579 | PR-27570 |
| BB4884 (PrTEF_DHCR7Ts) | *Tetraselmis* sp. GSL018 delta-7 sterol reductase codon optimised | PR-27581 | PR-27582 |
| BB4885 (PrTEF_DHCR7Wc) | *Waddlia chondrophila* delta-7 sterol reductase codon optimised | PR-27583 | PR-27584 |
| BB5041 (LoxP_HphMX) | pCfB5935, Holkenbrink *et al*, 2018<sup>33</sup> | PR-27972 | PR-27973 |
| BB5091 (*ERG4*_1kbUP_U1) | ST9100 gDNA | PR-27891 | PR-28358 |
| BB5092 (*ERG4*_1kbDOWN_U2) | ST9100 gDNA | PR-27893 | PR-27894 |
| BB5093 (*ERG5*_1kbUP_U1) | ST9100 gDNA | PR-27895 | PR-28359 |
| BB5094 (*ERG5*_1kbDOWN_U2) | ST9100 gDNA | PR-27897 | PR-27898 |
| BB5097 (*ERG4*_HphMX_KO) | BB5091 + BB5092 + BB5041 USER reaction | PR-27891 | PR-27894 |
| BB5098 (*ERG5*_HphMX_KO) | BB5093 + BB5094 + BB5041 USER reaction | PR-27895 | PR-27898 |
| BB5100 (TtSTC_USER) | *Tetrahymena thermophila* squalene-tetrahymanol cyclase codon optimised | PR-28361 | PR-28362 |
| BB5371 (PrGPAT_SMT2Cq) | *Chenopodium quinoa* sterol methyl transferase codon optimised | PR-27621 | PR-27786 |
| BB5373 (PrGPAT_DHCR24Sl) | *Solanum lycopersicum* delta-24(25) sterol reductase codon optimised | PR-27599 | PR-27764 |
| BB5456 (StSSR1_USER) | *Solanum tuberosum* delta-24(28) sterol reductase codon optimised | PR-29404 | PR-29405 |

**Supplementary Table 6.**
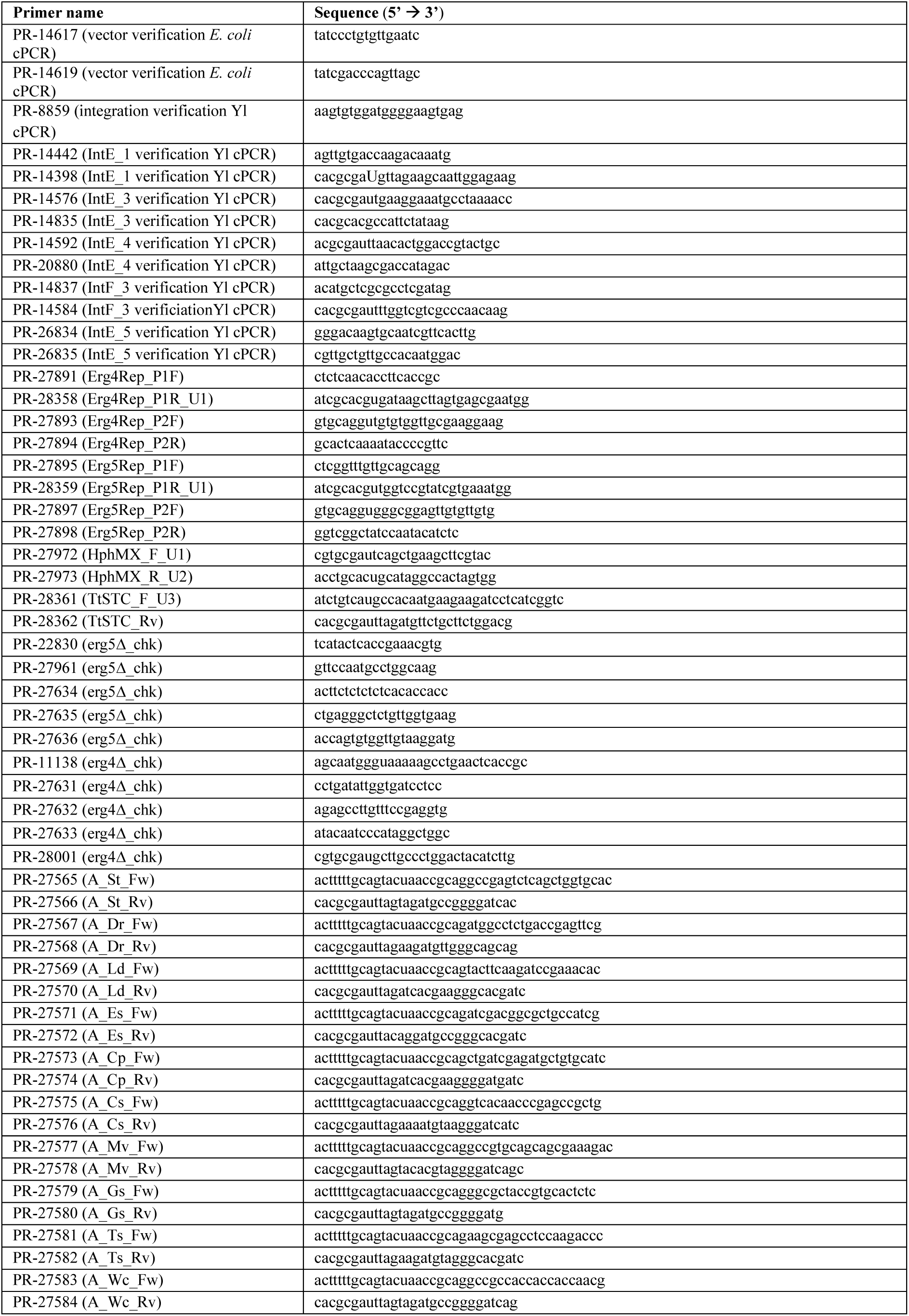
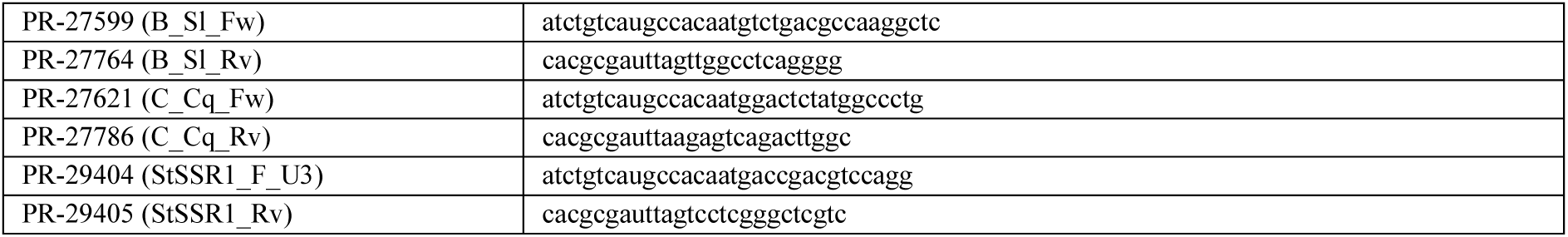
Primers used in this study.

**Supplementary Table 7.** Sterol content of diets prepared for feeding trials.

| Sterol | Dietary sterol content (w/w) |  |  |  | Percentage of total sterol (%) |  |  |  |
| --- | --- | --- | --- | --- | --- | --- | --- | --- |
|  | MxSt | WT | Tet | Base | MxSt | WT | Tet | Base |
| 24-Methylenecholesterol | 0.146 ± 0.040 | 0 | 0 | 0 | 43.3 ± 4.15 | 0.300 ± 0.068 | 0.228 ± 0.037 | 0.131 ± 0.051 |
| Campesterol | 0.038 ± 0.028 | 0.016 ± 0.000 | 0.015 ± 0.002 | 0.069 ± 0.011 | 13.5 ± 10.8 | 14.4 ± 1.28 | 11.3 ± 0.647 | 20.1 ± 1.45 |
| Cholesterol | 0.008 ± 0.000 | 0 | 0 | 0.001 ± 0.000 | 2.61 ± 0.496 | 0.421 ± 0.073 | 0.257 ± 0.022 | 0.209 ± 0.015 |
| Desmosterol | 0 | 0 | 0 | 0 | 0.114 ± 0.037 | 0 | 0 | 0 |
| Isofucosterol | 0.014 ± 0.005 | 0.016 ± 0.001 | 0.014 ± 0.001 | 0.053 ± 0.011 | 4.6 ± 2.38 | 14.2 ± 0.267 | 10.3 ± 0.876 | 15.9 ± 4.28 |
| Sitosterol | 0.022 ± 0.004 | 0.026 ± 0.002 | 0.023 ± 0.002 | 0.073 ± 0.004 | 7.06 ± 2.60 | 23.5 ± 1.42 | 16.6 ± 1.54 | 21.6 ± 2.45 |
| Tetrahymanol | 0.073 ± 0.054 | 0 | 0.037 ± 0.006 | 0 | 19.6 ± 13.3 | 0 | 27.1 ± 3.41 | 0 |
| Zymosterol | 0.001 ± 0.000 | 0.001 ± 0.000 | 0 | 0 | 0.220 ± 0.175 | 0.574 ± 0.118 | 0.200 ± 0.021 | 0 |
| Ergosterol | 0 | 0.015 ± 0.003 | 0.013 ± 0.001 | 0 | 0 | 13.3 ± 2.31 | 9.3 ± 0.427 | 0 |
| Campestanol | 0.002 ± 0.002 | 0.007 ± 0.000 | 0.007 ± 0.001 | 0.038 ± 0.008 | 0.422 ± 0.597 | 6.26 ± 0.168 | 5.1 ± 0.643 | 11.2 ± 2.15 |
| Cycloartenol | 0.004 ± 0.000 | 0.003 ± 0.000 | 0.003 ± 0.000 | 0.009 ± 0.001 | 1.15 ± 0.308 | 2.56 ± 0.421 | 2.04 ± 0.119 | 2.77 ± 0.398 |
| Stigmasterol | 0.005 ± 0.001 | 0.004 ± 0.000 | 0.003 ± 0.000 | 0.015 ± 0.003 | 1.53 ± 0.119 | 3.24 ± 0.294 | 2.36 ± 0.241 | 4.38 ± 0.501 |
| 31-Norcycloartanol | 0.005 ± 0.000 | 0.004 ± 0.001 | 0.004 ± 0.000 | 0.014 ± 0.002 | 1.47 ± 0.216 | 4.02 ± 0.261 | 3.25 ± 0.172 | 4.21 ± 0.484 |
| Sitostanol | 0.007 ± 0.002 | 0.013 ± 0.002 | 0.010 ± 0.001 | 0.044 ± 0.034 | 2.17 ± 0.390 | 11.3 ± 0.794 | 7.61 ± 1.33 | 11.8 ± 8.11 |
| 24(28)-Methylenecycloartanol | 0.007 ± 0.000 | 0.007 ± 0.001 | 0.006 ± 0.001 | 0.026 ± 0.005 | 2.22 ± 0.582 | 5.93 ± 0.453 | 4.35 ± 0.127 | 7.69 ± 2.10 |
| Total | 0.331 ± 0.061 | 0.111 ± 0.007 | 0.136 ± 0.010 | 0.341 ± 0.045 | 100 | 100 | 100 | 100 |

## Supplementary Data 1. (separate file)

Zu *et al.* (2021) Pollen sterol data bee-pollinated subset.

## Supplementary Data 2. (separate file)

Colony sample GCMS statistical analysis.

## Supplementary Data 3. (separate file)

Hive temperature and humidity recordings.

## Acknowledgements

Acknowledgements: We thank Prof David Nes and Wenxu Zu for initial discussions about engineering yeast for sterols, Paulina Rubaszka for support with Ambr® 250 bioreactors, Ann Fink Jensen for help with bioreactor autoclavation and Christiane Glitz for assistance with HPLC. We thank Daniel Stabler and Jennifer Chennells for preliminary feeding trials, Harry Bridge and Lea Beaupere for beekeeping assistance during feeding trials, Samuel Furse for help with freeze-drying, and Lea Beaupere and Kieran Walter for data transcription. Temperature and humidity sensors were provided by BeeHero. EM was funded by BBSRC iCASE grant (BB/M011224/1) with Apix Nutrition as an industrial partner. This work was funded in part by BBSRC grant BB/T015292/1 awarded to GAW.

## Contributions

EM, GAW and IB conceptualised the study. EM, GAW and IB designed yeast experiments; EM performed yeast experiments. EM, RTdS, and GAW designed honeybee feeding trials; RFSG, RTdS, and GAW developed the honeybee feeding assay; EM and RTdS carried out honeybee feeding trials. JDD, EM and JAA performed heterologous gene searches. DF and PS performed mass spectrometry and identified sterols. SF provided and analysed temperature and humidity sensor data. EM conducted data analysis. EM and GAW wrote the manuscript. All authors commented on the article and approved the submitted version.

